# The modified activity of prolyl 4 hydroxylases (P4Hs) reveals the effect of arabinogalactan proteins (AGPs) on changes in the cell wall during the tomato ripening process

**DOI:** 10.1101/2024.01.22.576594

**Authors:** Nataliia Kutyrieva-Nowak, Agata Leszczuk, Lamia Ezzat, Dimitris Kaloudas, Adrian Zając, Monika Szymańska-Chargot, Tomasz Skrzypek, Afroditi Krokida, Khansa Mekkaoui, Evangelia Lampropoulou, Panagiotis Kalaitzis, Artur Zdunek

## Abstract

Arabinogalactan proteins (AGPs) are proteoglycans with an unusual molecular structure characterised by the presence of a protein part and carbohydrate chains. Their specific properties at different stages of the fruit ripening programme make AGPs unique markers of this process. An important function of AGPs is to co-form an amorphous extracellular matrix in the cell wall-plasma membrane continuum; thus, changes in the structure of these molecules can determine the presence and distribution of other components. The aim of the current work was to characterise the molecular structure and localisation of AGPs during the fruit ripening process in transgenic lines with silencing and overexpression of *SlP4H3* genes. The objective was accomplished through comprehensive and comparative *in situ* and *ex situ* analyses of AGPs from the fruit of transgenic lines and wild-type plants at specific stages of ripening. The experiment showed that changes in P4H3 activity affected the content of AGPs and the progress in their modifications in the ongoing ripening process. The analysis of the transgenic lines confirmed the presence of AGPs with high molecular weights (120–60 kDa) at all the examined stages, but a changed pattern of the molecular features of AGPs was found in the last ripening stages, compared to WT. In addition to the AGP molecular changes, morphological modifications of fruit tissue and alterations in the spatio-temporal pattern of AGP distribution at the subcellular level were detected in the transgenic lines with the progression of the ripening process. The work highlights the irreversible impact of AGPs and their alterations on the fruit cell wall assembly and changes in AGPs associated with the progression of the ripening process.

**GRAPHICAL ABSTRACT:** 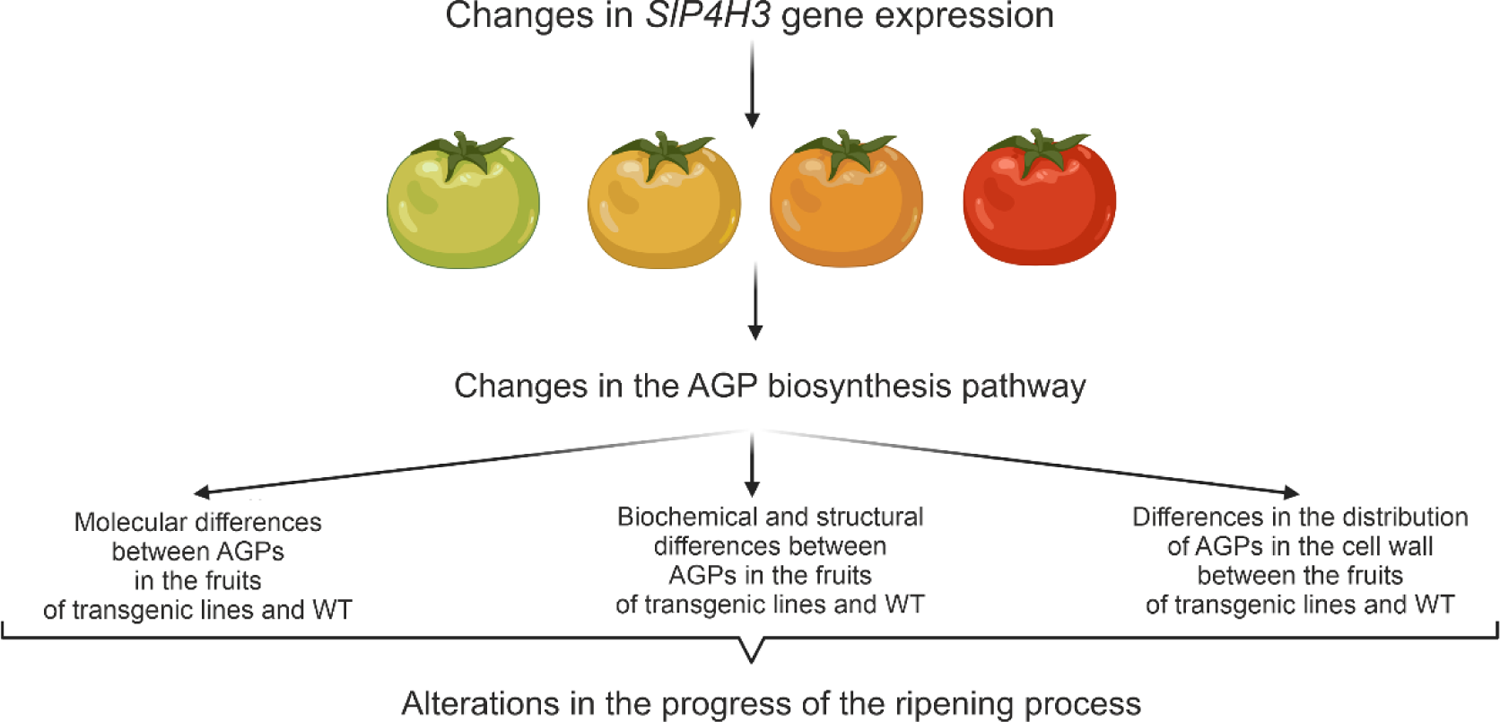

## 1. INTRODUCTION

### 1.1. Arabinogalactan proteins (AGPs) in the fruit ripening process

The tomato (*Solanum lycopersicum* L.) is one of the most widely cultivated crops worldwide and a well-studied model for tissue-specific gene modification and fruit ripening (Feder et al., 2020). In turn, studies of fruit ripening are useful for both practical agricultural applications and improvement of the understanding of physiological programmes. The ripening process is a complex phenomenon inducing changes in fruit tissues. Alterations in fruit texture and colour are attributed to different metabolic processes in the cell wall (Orfila et al., 2002). Our previous studies have shown that there are substantial changes in the structure and distribution of arabinogalactan proteins (AGPs) correlated with the ongoing ripening process (Kutyrieva-Nowak et al., 2023a). As common components in the plant extracellular matrix, AGPs are attached to the plasma membrane via glycosylphosphatidylinositol (GPI) anchors in the C-terminal domain (Showalter, 2001; Lopez-Hernandez et al., 2020; Zhou, 2022; Leszczuk et al., 2023). The specific and mutable distribution of AGPs in the cell wall-plasma membrane continuum allows identification of the effects of these molecules on the specific arrangement of the cell wall assembly in fruit cells (Liu et al., 2015; Leszczuk et al., 2018). Moreover, in the newest concept of the cell wall model named the APAP1 complex, AGPs are considered as cross-linkers between other cell wall components in which they are covalently attached to other polysaccharides of the cell wall, such as hemicellulose and pectins (Tan et al., 2013; Hijazi et al., 2014a, 2014b; Leszczuk et al., 2020c). Furthermore, it should also be underlined that some processes of plant growth and development, including cell expansion, somatic embryogenesis, root and stem growth, signalling during cell-cell communication, salt tolerance, and programmed cell death, require the presence of AGPs (Showalter, 2001; Zhang et al., 2020; Leszczuk et al., 2023). Although the involvement of AGPs in many aspects of plant growth and development is well understood, still little is known about their function in fruit development and ripening. It is well-known that hydroxyproline-rich glycoproteins (HRGPs), i.e. the group to which AGPs are classified, are involved in fruit softening and affect the progression of the ripening process (Fragkostefanakis et al., 2012). Moreover, our previous investigations conducted on apple fruit after colonisation of *Penicillium spinulosum* have shown that the amount of AGPs increased in infection-associated modifications in fruit tissue. The increased amount of AGPs during the development of fungal disease is correlated with their assumed contribution in response to biotic stress, i.e. structure of the mechanical barrier against plant pathogens (Leszczuk et al., 2019c, 2020b).

AGPs are one of the most complex protein families in the plant cell wall. Notably, AGPs contain carbohydrate chains in 90-95% of their molecular mass and the protein core constitutes only 5-10%. Among others, AGPs have unique characteristics, such as the presence of arabinogalactan type II (AG) chains attached to the protein core composed of hydroxyproline (Hyp) residues and typical Ala-Pro, Pro-Ala, Thr-Pro, Ser-Pro, Val-Pro, and Gly-Pro peptide repeats (Showalter, 2001; Liu et al., 2015; Leszczuk et al., 2020b). In turn, the β-1,3-galactose chains of AGPs are modified by linking β-1,6-galactose side chains, which are further modified by the addition of arabinose (Ara), rhamnose (Rha), fucose (Fuc), xylose (Xyl), and β-glucuronic acid (GlcA) residues. The synthesis of the AGP molecule occurs in several steps, including the addition and removal of amino acids and carbohydrate residues; therefore, changes in any of these steps may result in structural and functional modifications of AGPs (Fragkostefanakis et al., 2012; Leszczuk et al., 2023). During the AGP synthesis pathway, an important function is performed by prolyl 4 hydroxylases (P4Hs). After proper hydroxylation of proline catalysed by P4Hs, AGPs undergo glycosylation, and this molecular mechanism is well known, but the effect of changes in this mechanism on AGPs as well as the mechanism of disruption of the subsequent glycosylation process by changes in the protein domain need to be elucidated.

### 1.2. Modification of AGP carbohydrate chains

It is assumed that the wide functionality of AGPs is the result of their complex glycome structure (Showalter, 2001). In more detail, it is suggested that GlcA may confer functional properties on AGPs specifically linked to β-(1→6)-galactan chains (Tan et al., 2004; Leszczuk et al., 2023). One of the presumed mechanisms of the alterations in the activity and distribution of AGPs is their interaction with Ca^2+^, which depends on the bonds between GlcA residues and Ca^2+^ (Lamport and Várnai, 2013). As reported by Lamport and Várnai, (2013) glucuronidated AGPs may interact with Ca^2+^ and then release Ca^2+^. In β-glucuronyltransferase mutants (*GlcAT)*, a reduction of Ca^2+^ binding with AGPs, which affected the cell wall and plasma membrane stabilisation and caused less intensive plant growth, was revealed. Binding and release of Ca^2+^ by AGPs may contribute to the regulation of cellular signalling (Lopez-Hernandez et al., 2020), and AGPs may serve as a reservoir of Ca^2+^ affecting plant development (Lamport et al., 2018). In our study on apple fruit, the quantitative analysis of the Ca^2+^ content in extracted AGPs showed that the amount of Ca^2+^ increased with the progression of the ripening process (Leszczuk et al., 2020a). The maximum Ca^2+^ content in AGPs determined in red mature fruit was eight-fold higher than in AGPs extracted from unmatured green fruit (Leszczuk et al., 2020a). Interestingly, the enhancement of Ca^2+^ accumulation by AGPs in the red fruit was correlated with an increase in the content of GlcA, which participates in the formation of complexes with Ca^2+^ (Lamport and Várnai, 2013; Leszczuk et al., 2020a). Furthermore, studies performed by Lopez-Hernandez and coworkers (2020) also revealed an interaction between AG chains and Ca^2+^ and showed that AGPs provided a source of apoplastic Ca^2+^. AGs from mutant plants exhibited reduced glucuronidation, which caused binding of less Ca^2+^ *in vitro* and development deficiencies in these plants.

The AGPs from *GlcAT14a/b/d-1* mutants bound nearly 80% less Ca^2+^ than AGPs from wild-type plants (WT) (Lopez-Hernandez et al., 2020). Interestingly, the research carried out by Nibbering and coworkers (2020) has identified and characterised the enzymatic activity of two exo-β-1,3-galactosidases (GH43) whose activity influenced the levels of AGPs in the cell wall. The analysis of quantification of cell wall-associated AGPs has confirmed that the *Gh43null* mutant differs from WT, which may clarify the GH43 role in the regulation of the AGP cell wall affinity. The aforementioned investigations showed that GlcA also forms a bridging residue between pectin and AGPs, confirming the presence of the APAP1 complex (Tan et al., 2013, 2023). Moreover, it can be concluded that APAP1 may be the target of GH43 activity, as it affects cell wall-bound AGPs (Nibbering et al., 2020).

### 1.3. Modification of the protein moiety of AGPs – effect of prolyl 4 hydroxylases (P4Hs)

As mentioned above, P4Hs catalyse proline hydroxylation, a major post-translational modification of HRGPs (Kivirikko and Myllyharju, 1998; Hieta and Myllyharju, 2002; Koski et al., 2009; Leszczuk et al., 2020c, 2023). The impact of P4Hs in plant growth and development was demonstrated in carrot root *in vivo* (Cooper and Varner, 1983). Structural alterations in HRGPs were correlated with abnormal cell division, which influenced the loosening of the cell wall matrix and the *de novo* synthesis and rearrangement of cell wall components (Cooper et al., 1994). In other studies, mutants of *Arabidopsis* with T-DNA knockout P4Hs had shorter root hair, which was caused by destruction of extensins, and were characterised by a disrupted *O*-glycosylation process (Velasquez et al., 2015). Overall, in plants with disturbed function of P4Hs, the total hydroxyproline content is changed, leading to structural and functional changes during cell division and expansion (Vlad et al., 2007; Perrakis et al., 2019; Konkina et al., 2021). Also, alterations in tomato P4H genes caused by virus-induced gene silencing (VIGS) allowed determination of the impact on the weight and size of leaves resulting in intensified biomass production (Fragkostefanakis et al., 2014).

Despite the well-documented importance of HRGPs, there is still insufficient information on the impact of P4H activity on AGPs. Phenotypic analyses have confirmed that overexpression and silencing of P4H genes cause changes in the *O*-glycosylation process, and the role of AGPs in the fruit ripening process may be determined via genetic manipulation of the protein backbone which changes glycan moieties. The purpose of our study was to characterise the molecular structure and distribution of AGPs during the fruit ripening process in transgenic lines with silencing and overexpression of P4H genes. The aims were accomplished through *in situ* and *ex situ* analyses as well as a comparative analysis of AGPs in the fruit of transgenic lines and WT at different stages of ripening. The development of transgenic lines with altered *SlP4H3* gene expression facilitated the determination of modification in the structure and arrangement of AGPs in the cell wall, thus contributing to the elucidation of the effect of these molecules on the whole extracellular matrix.

## 2. MATERIAL & METHODS

### 2.1. The SlP4H3 RNAi and overexpression transgenic lines

Homozygous T2 and T3 generation tomato (*Solanum lycopersicum* cv. ‘Ailsa Craig’) plants of two RNAi, RNA#1, RNA#7, and four overexpression, OEX#1, OEX#2, and cGFP OEX (cGFP), nGFP OEX (nGFP), lines were grown in the glasshouses at Chania, Greece under standard conditions (Perrakis et al., 2021). Tomato fruits were harvested at the Breaker (BR), Turning (TU), Pink (PINK), and Red Ripe (RR) previously described by Nakatsuka and coworkers (1998). The ripening stages were identified by assessment of changes in the fruit colour as proposed by Batu (2004). The first stage of the ripening process is BR. It is the stage after development, in which tomatoes have a pale-green surface and the first signs of ripening (an orange colour) are visible. The next stage is TU, in which the first appearance of a pale-pink colour of the fruit surface is evident on 10-30% of the surface. The next stage is PINK, in which two-thirds (above 60%) of the fruit surface is red. The last stage is RR, in which the fruit is red on the entire surface (Nakatsuka et al., 1998; Kutyrieva-Nowak et al., 2023a)

### 2.2. RNA extraction and cDNA synthesis

Total RNA was extracted from 200 mg of pericarp tissue at the fruit ripening stages of BR, TU, PK, and RR from the RNA#1, RNA#7, OEX#1, OEX#2, cGFP, nGFP lines and wild-type (WT). The samples were pulverized in liquid nitrogen and RNA was extracted using the NucleoZOL reagent (MACHEREY-NAGEL, Germany), following the manufacturer’s instructions. The RNA samples were treated with DNase I (NEB, RNase-free, Ipswich, MA, USA) and approximately 1 μg of RNA was reverse transcribed using SuperScriptTM II RT (Invitrogen, Carlsbad, CA, USA) for cDNA synthesis following the manufacturer’s instructions.

### 2.3. Real-time qRT-PCR analysis

Real-time PCR analysis was conducted using the SYBR™ Select Master Mix (Thermo Fisher Scientific, Waltham, MA, USA) in a CFX Connect™ Real-Time PCR Detection System (Bio-Rad, Hercules, CA, USA). The cDNA samples were normalized with b-actin as the reference gene using the primers SlActinF 5’-GTCCCTATTTACGAGGGTTATGCT-3’, SlActinR 5’-GTTCAGCAGTGGTGGTGAACA-3’. The primers used to determine the SlP4H3 expression in the RNA#1, RNA#7, OEX#1, OEX#2, cGFP, nGFP lines and WT were SlP4H3 Forward 5’-GTGAAAGGAAGGCATTCTCG-3’ and SlP4H3 Reverse 5’-CTTTCTGAGAGC CCCTGTGA-3’. The PCR conditions started at 50 °C for 2 minutes and 95 °C for 5 min, followed by 40 cycles of denaturation at 95 °C for 15 s, annealing at 60 °C for 30 s, and extension at 72 °C for 30 s. A final melt-curve stage was performed, with temperatures set at 95°C for 15 seconds, followed by 60°C for 30 seconds with a 0.5°C increment at each repeat. Data analysis was carried out using the 2-ΔΔCT method (Livak and Schmittgen, 2001), and the results were presented as relative levels of gene expression with actin as the internal standard. Standard errors were calculated for all mean values. Three biological replicates were used per sample.

### 2.4. Molecular analyses

#### 2.4.1. Protein extraction

Protein extraction was conducted according to Ling’s protocol (Ling et al., 2021) and with modifications described in our previous work (Kutyrieva-Nowak et al., 2023a). The fruit tissue was cut into cube-shaped pieces and frozen at −80°C before the extraction procedure. The tomato fruit tissue was homogenised to a fine powder in liquid nitrogen, and then Laemmli’s extraction buffer (1970) was added. The modified Laemmli’s buffer contained 65mM Tris-HCl pH 6.8, 2% SDS, 2mM EDTA, 1mM PMSF, 700mM β-mercaptoethanol, and a 1:10 protease inhibitor. The samples were boiled at 95°C for 5 minutes and then clarified by centrifugation at 14 000 rpm at 4°C for 20 minutes. The final step was to collect the supernatant and freeze it at −80°C for the next assays.

#### 2.4.2. Dot blotting with quantitative analysis

For determination of the presence of AGPs in the supernatants, an immuno-dot-blot reaction using specific antibodies against the carbohydrate epitopes of AGPs was carried out. The commercial monoclonal antibodies used to study AGPs were provided by Kerafast (USA). The experiment was conducted using primary antibodies: JIM13, which recognises the trisaccharide epitope β-D-GlcA-(1,3)-α-D-GalA(1,2)-α-L-Rha (Knox et al., 1991; Yates and Knox, 1994; Yates et al., 1996), LM2, which recognises the epitope β-linked GlcA (Yates and Knox, 1994; Smallwood et al., 1996; Yan et al., 2015), LM14, which recognises the epitope arabinogalactan type II (Moller et al., 2008), and LM1, which recognises extensin glycoprotein (Smallwood et al., 1996). Each sample was dotted onto a pre-prepared nitrocellulose membrane (PVDF) with a 0.2 µm pore size (Thermo Scientific, USA) and blocked using a 5% solution of bovine serum albumin (BSA; Sigma, USA) in phosphate-buffered saline (PBS; Sigma, USA). After washing in Tris-buffered saline (TBST, 7.6 pH) three times, the membranes were incubated for 2 h at room temperature (RT) with primary antibodies diluted 1:500 in 2.5% BSA in PBS. After washing with TBST, the membranes were incubated for 2 h at RT with secondary antibodies Anti-Rat-IgG conjugated with alkaline phosphatase (AP) (Sigma-Aldrich, USA) at a dilution of 1:1000 in 2.5% BSA in PBS. After three washing steps in TBST and AP-buffer, AGPs were finally detected using the following substrates: 4 mg of 5-bromo-4-chloro-3-indolylphosphate (BCiP; Sigma, USA) in 1 mL water and 9 mg of nitro-blue tetrazolium (NBT; Sigma, USA) in 0.3 mL water and 0.7 mL N, N-dimethylformamide (DMF; Thermo Scientific, USA) in the dark. After membrane imaging using GelDoc Go Imaging System (Bio-Rad, USA), the measurement of the colour intensity of the dots was carried out using Image Lab Software v. 6.1 (Bio-Rad, USA). The heat map was prepared using Microsoft tools in which colour intensity is proportional to a numerical value. The numerical values are rearranged to ×10^3^ for better representation of results. Data for each sample were obtained from three independent experiments.

#### 2.4.3. SDS-PAGE and Western blotting with quantification

Gel electrophoresis and next Western blotting were performed for identification of the molecular weight of AGPs from the fruit tissue in the different stages of ripening. After total protein extraction, quantification of the protein content was carried out using the Bradford assay. Protein separation was performed using SDS-PAGE with 12.5% resolving gel and 4% stacking gel. After SDS-PAGE electrophoresis, proteins with the gel were electroblotted onto a PVDF membrane. After Western blotting wet transfer, the membrane was washed in TBST on a shaking platform. After washing, the preincubation step was performed for 1 h at RT with 5% BSA in PBS, and the membrane was incubated for 2 h at RT with primary antibodies at a concentration of 1:500 in 2.5% BSA in PBS. After washing with TBST, the membrane was incubated for 2 h at RT with secondary antibodies conjugated with AP at a concentration of 1:1000. After washing with TBST and AP-buffer three times, visualisation of bands was performed using BCiP and NBT at a concentration described above in the GelDoc Go Imaging System (Bio-Rad, USA). For qualitative and quantitative analysis, the Pierce™ Prestained Protein MW Marker (Thermo Scientific, USA) was used. The band thickness, width, and colour depth were used for qualitative analysis. The quantitative analysis of protein bands was performed by measurements of obtained stripes using Image Lab Software v. 6.1 (Bio-Rad, USA). Three independent analyses were carried out.

#### 2.4.4. ELISA with quantitative analysis

Based on the number of antigen-antibody interactions produced, ELISA is a fundamental test for both qualitative and quantitative identification of AGPs in samples (Pfeifer et al., 2020; Kutyrieva-Nowak et al., 2023b). First, the samples were added to each well on a 96-well plate (Nunc MaxiSorpTM flat-bottom, Thermo Fisher Scientific, Denmark) and immobilized at 37°C for 72 h with shaking (350 rpm). Then, the coated plate was washed three times with PBS (pH 7.4), preincubated for 1 h at 37°C with 0.1% BSA in PBS, and incubated for 1 h at 37°C with primary antibodies at a concentration of 1:20 in PBS. After the washing step, the plate was incubated for 1 h at 37°C with the secondary antibody (in a dilution of 1:500 in PBS). After the incubation and washing steps, the reaction was developed by adding a freshly prepared substrate solution of p-nitrophenol phosphate (PNPP) according to Thermo Scientific instructions. The reaction was run in the dark and stopped with 2M NaOH after 20 minutes. The absorbance was measured in an ELISA reader (MPP-96 Photometer, Biosan) at 405 nm and analysed with Statistica v.13 tools (TIBCO Software Inc. USA). Analysis of variance (one-way ANOVA) and Tukey’s Honestly Significant Difference (HSD) post hoc test were used to compare the mean results. For all analyses, the significance level was estimated at p < 0.05.

### 2.5. Biochemical–structural analyses

#### 2.5.1. Extraction of AGPs using β-glucosyl Yariv Reagent

AGPs were isolated from tomato fruit tissue using the β-glucosyl Yariv Reagent (β-GlcY; Biosupplies, Australia) according to the extraction protocol proposed by Lamport (2013) and used in our previous studies (Leszczuk et al., 2020a, 2020b; Kutyrieva-Nowak et al., 2023a). The liquid-nitrogen pre-frozen and homogenised fruit tissue was added to 2% CaCl_2_ and incubated at RT. After 3 h, the homogenate was centrifuged at 10 000 rpm for 30 minutes at RT. The supernatant was retained and added to 1 mg/mL β-GlcY in 2% CaCl_2_ and left overnight at RT. The insoluble Yariv-AGP complex was centrifuged for 15 minutes at 2 000 rpm, and the precipitate was resuspended in MiliQ water. Next, sodium metabisulphite (Thermo Scientific, USA) was gradually added to the precipitate and heated at 50°C to reduce the diazo linkage until the suspension was decolourised. After obtaining a clear yellow solution, the suspension was transferred to dialysis tubing with a 12-kDa MW cut-off (32 mm flat width; Sigma, USA) and stirred overnight. The dialysate was lyophilized, and then the AGPs were weighed.

#### 2.5.2. FTIR analyses

FTIR spectra were collected on a Nicolet 6700 FTIR spectrometer (Thermo Scientific, Madison, WI, USA). The Smart iTR ATR sampling accessory was used. The spectra of the lyophilized AGP extracts were collected over the range of 4000–650 cm^−1^. Three samples of each material were examined in the same conditions. For each sample, 200 scans were averaged with a spectral resolution of 4 cm^−1^. Then, a final average spectrum was calculated for a given material. These spectra were baseline corrected and normalized to 1.0 at 1019 cm^−1^. To highlight the differences between the samples, the principal component analysis (PCA) of the spectra was performed in the mid-infrared region of 4000–400 cm^−1^ using Unscrambler 10.1 (Camo Software AS., Norway). The maximum number of components taken for the analysis was five, and the NIPALS algorithm was used (Szymanska-Chargot et al., 2015; Chylińska et al., 2016). The visualisation of FTIR spectra and PCA analysis were performed using the OriginPro program (Origin Lab v. 8.5 Pro, Northampton, USA).

### 2.6. Microscopic analyses

#### 2.6.1. SEM imaging

For morphological description, AGPs extracted using β-glucosyl Yariv Reagent from fruit tissue at different stages of the ripening process (the BR and RR stages) were imaged with a scanning electron microscope (SEM, Zeiss Ultra Plus, Oberkochen, Germany) in high vacuum (5 × 10^−3^ Pa) using a secondary electron detector at 3 kV.

#### 2.6.2. Preparation of material for immunocytochemistry

The microscopic analyses were carried out in accordance with our previous papers (Leszczuk et al., 2019a; Kutyrieva-Nowak et al., 2023a, 2023b). For the microscopic examination, the fruit tissue was subjected to the procedure of fixation, resin embedding, and sectioning. Cube-shaped pieces of the fruit tissue were fixed in 2% paraformaldehyde (Sigma, USA) and 2.5% glutaraldehyde (Sigma, USA) in PBS and placed in vacuum (0.7 bar) seven times for 10 minutes each and then left overnight. Next, the samples were washed in PBS and distilled water and dehydrated in graded series of ethanol solutions (30%, 50%, 70%, 90%, and 96% for 15 minutes each and 99.8% twice for 30 minutes). The tissue was placed in 99.8% ethanol and LR White resin (at a ratio of 3:1, 1:1, 1:3; Sigma Aldrich, USA), and next in 100% resin LR White. The polymerisation was carried out for 48 h at 55°C. For CLSM imaging, the samples were cut into semi-thin sections (1 µm) using a glass knife-equipped ultramicrotome (PowerTome XL, RMC Boeckeler, USA). Next, the sections were placed on poly-L-lysine coated glass slides (Sigma, USA). For TEM imaging, the samples were cut into ultra-thin sections (70 nm) using a diamond knife-equipped ultramicrotome, and the sections were placed on formvar film-coated nickel square grids (EM Resolutions Ltd, UK).

#### 2.6.3. Immunofluorescence method with confocal laser scanning microscopy (CLSM imaging)

Immunofluorescence imaging of AGP epitopes at the tissue/cellular level facilitates evaluation of changes in the distribution of AGPs in fruit at different ripening stages. The experiment was conducted using JIM13, LM2, and LM14 antibodies. Semi-thin sections on poly-L-lysine coated glass slides were circled with a liquid blocker Dako-Pen (Sigma Aldrich, USA). The samples were washed and pre-incubated for 30 minutes at RT with 2% BSA in PBS to block non-specific binding sites. After the washing step, the sections were incubated with primary antibody diluted 1:50 in 0.1% BSA in PBS overnight at 4°C. After washing with PBS three times, secondary Alexa Fluor 488 antibodies (diluted 1:200 in 0.1% BSA in PBS, goat Anti-Rat-IgM, Thermo Fisher Scientific, Denmark) were added and the sections were incubated overnight at 4°C. Then, the incubated sections were washed in PBS and MiliQ water and finally counterstained with Calcofluor White (Sigma Aldrich, USA). An Olympus BX51 CLSM microscope with corresponding software FluoView v. 5.0. (Olympus Corporation, Tokyo, Japan) was used for imaging. In order to perform control reactions, the incubation with the primary antibody was omitted. All photographs, figures, and schemes were edited using the CorelDrawX6 graphics program.

#### 2.6.4. Immunogold labelling technique with the transmission electron microscope (TEM imaging) and quantitative analysis

Immunogold labelling of AGP epitopes facilitates identification and visualization of changes in their distribution at the subcellular level. The experiment was conducted using JIM13, which recognises the most common and specific AGP epitope in fruits. Ultra-thin sections placed on grids were washed in distilled water, and next pre-incubated for 30 minutes at RT with 1% BSA in PBS. Then, the grids were incubated for 3 h at 37°C with the primary antibody (diluted 1:10 in 0.1% BSA in PBS), washed in the blocking solution, and labelled with the secondary antibody (diluted 1:50 in 0.1% BSA in PBS, Anti-Rat-IgG – Gold antibody; Sigma, USA) for 1 h at 37°C. After washing in PBS and distilled water, the samples were stained with a 1% uracyl acetate solution (for 10 minutes) and Reynold’s reagent (for 7 minutes). The observations were carried out using a transmission electron microscope (TEM Zeiss EM900) operating at 80 kV acceleration voltage (Carl Zeiss AG, Oberkochen, Germany) and equipped with a digital camera with corresponding software ImageSP v. 1.1.2.5. In the quantitative analysis of AGP labelling density, gold particles on the same size (2048 × 2048 pixels) micrographs were manually identified and counted with the software ImageJ v. 1.51. The results were expressed as the number of gold particles per 1 μm^2^ area of the examined cell compartment (Corral-Martínez et al., 2016; Leszczuk et al., 2018). All photographs, figures, and schemes were edited using the CorelDrawX6 graphics program.

## 3. RESULTS

### 3.1. Expression of SlP4H3 at fruit ripening stages in overexpression and silencing lines

Two SlP4H3 RNAi independent lines, #1 and #7, and two SlP4H3 overexpression lines, OEX#1, OEX#2 were previously produced (Perrakis et al., 2019, 2021). Moreover, two additional, independent overexpression lines were created, cGFP OEX (cGFP) and nGFP OEX (nGFP), comprising a construct in which a GFP tag was either at the C- or the N-terminus of the *SlP4H3* cDNA under the control of the 35S promoter (Perrakis et al., 2021). The expression levels of SlP4H3 in the two silencing and the four overexpression lines were determined at the four ripening stages (Figure 1). Lower expression levels were observed by approximately 50% at the RNAi#1 and #7 fruits in all ripening stages (Figure 1). In the overexpression lines, the highest levels of expression were detected in the cGFP and nGFP lines up to approximately 92- and 20-fold, respectively (Figure 1). In the OEX#1 and #2 lines, the *SlP4H3* transcript abundance was higher up to 4- and 8-fold, respectively (Figure 1).

**Figure 1.**
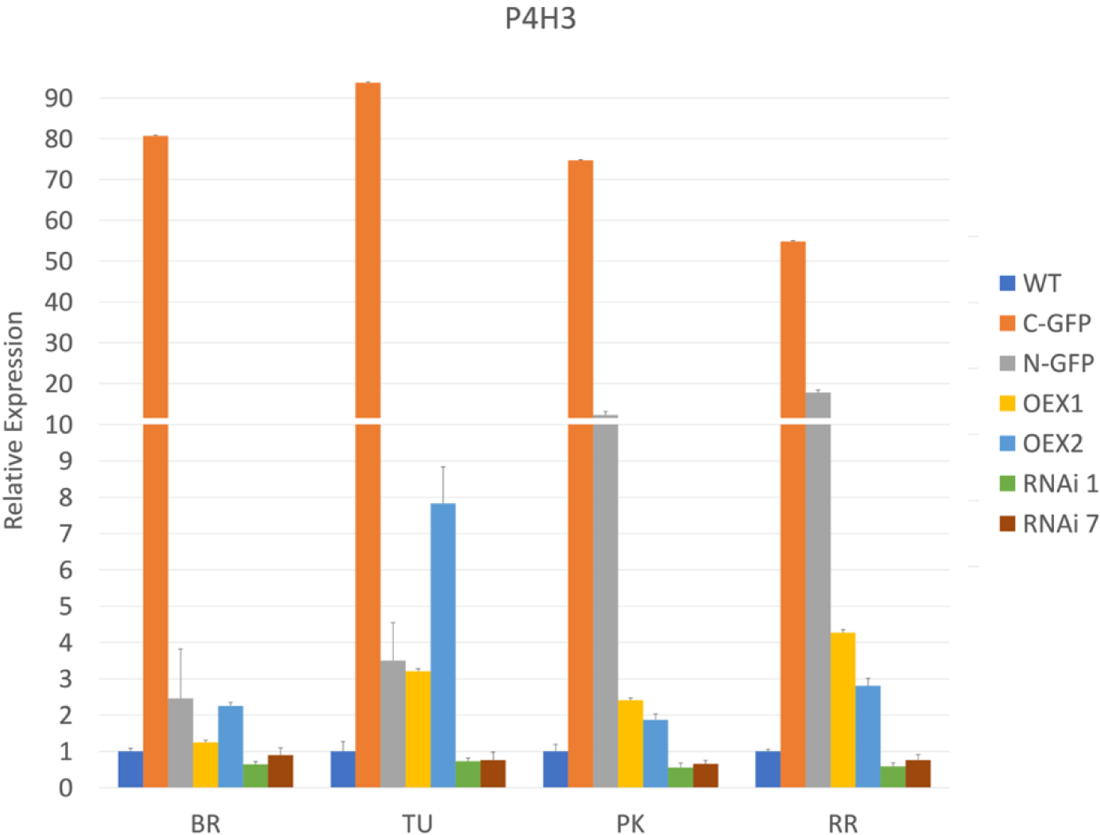
Expression of SlP4H3 at Breaker (BR), Turning (TU), Pink (PK) and Red Ripe (RR) fruit ripening stages in cGFP OEX, nGFP OEX, OEX #1, #2 and RNA #1, #7 lines and WT. The relative expression was calculated based on the comparative CT method and actin was used as the internal standard. The relative fold differences for each sample were determined by normalizing the Ct value for the *SlP4H3* gene to the Ct value for actin and calculated using the formula 2^-ΔΔCt^. Three biological replicates were performed. Error bars represent standard error.

### 3.2. Molecular differences

The molecular differences between AGPs in the fruit of the transgenic lines and WT at different stages of ripening were evaluated in several steps. Firstly, an immuno-dot-blot assay was performed to quickly check the presence of AGP epitopes. The Western blotting method helped to determine the proportions of the molecular weights of the AGP epitopes in the fruit samples. Afterwards, selective glycome profiling of AGPs using the enzyme-linked immunosorbent assay (ELISA test) was conducted to visualise quantitative changes in the AGP content in the fruit of the examined transgenic lines during the ongoing ripening process.

#### 3.2.1. Semi-quantitative determination of the sample composition

To validate the protein extraction and check the presence of specific AGP epitopes in the samples, an immuno-dot-blot assay was performed (Figure 2) and monoclonal antibodies against AGP epitopes (JIM13, LM2, and LM14) and against extensin (LM1) were used. The heat map obtained showed that the examined epitopes were present in different concentrations depending on the stage of the ripening process and the modification of P4H3 activity caused changes in the content of particular epitopes in the tested samples.

**Figure 2.**
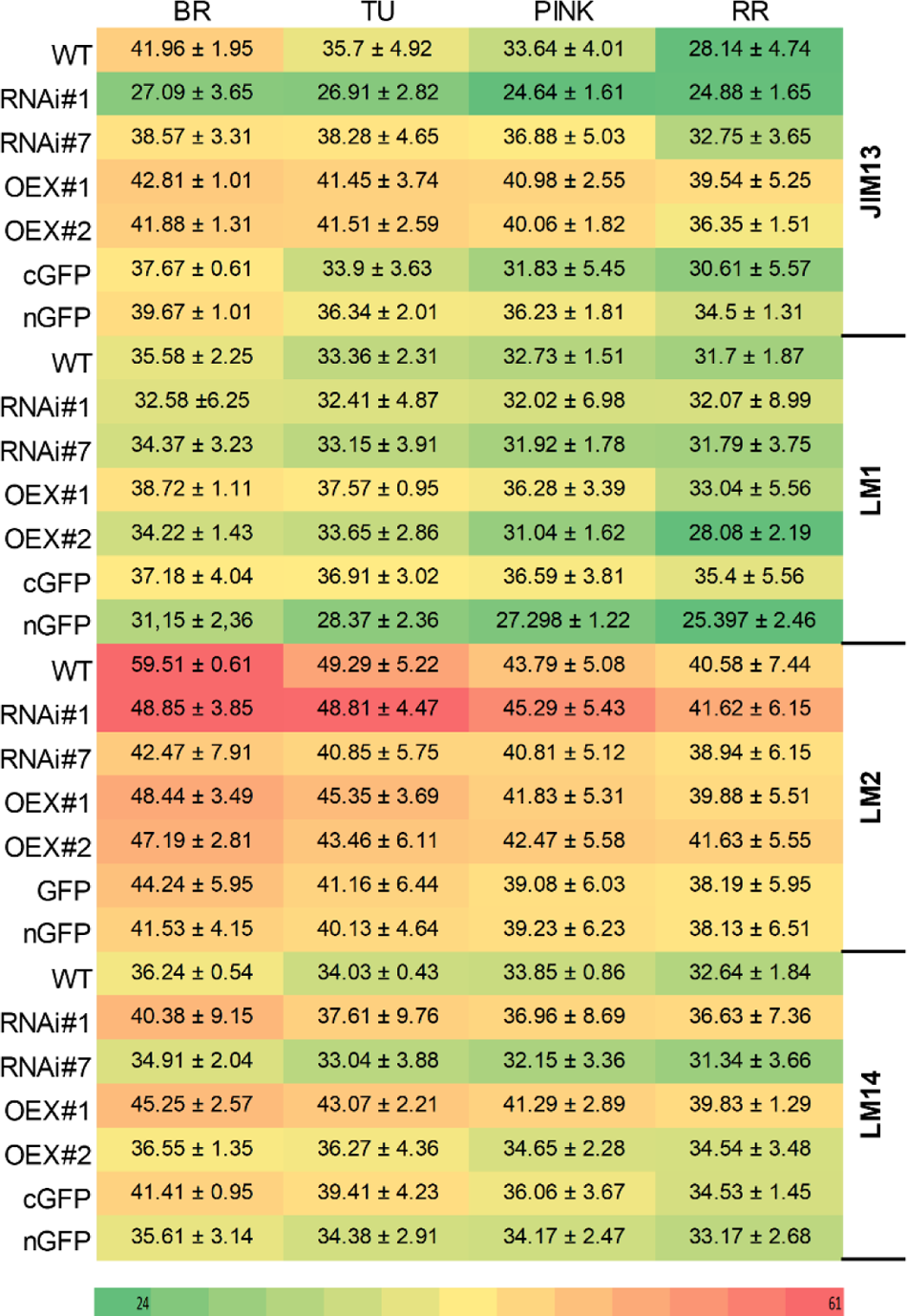
Examination of samples using JIM13, LM1, LM2 and LM14 antibodies at different ripening stages from the fruit of wild-type plants (WT) and transgenic lines: RNAi#1, RNAi#7, OEX#1, OEX#2, cGFP, and nGFP. The numerical value in the heatmap is ×10^3^. Abbreviations: BR – Breaker stage, TU – Turning stage, PINK – Pink stage, RR – Red Ripe stage

First of all, when analysing the ripening process itself, it is clear that the highest amount of all examined epitopes was detected at the BR stage, regardless of the line being tested. Additionally, in all cases, the number of all epitopes gradually decreased with the progression of the ripening process.

Secondly, the analysis of individual epitopes makes it possible to find differences in their content between individual lines. The assay of the fruit tissue of the RNAi#7 line with all the antibodies used indicated that silencing of the *SlP4H3* gene resulted in reduced intensity of the dots, compared to WT. In the same way, the analysis of the overexpression lines also revealed increased content of the tested epitopes in comparison to WT. The best visualisation effect of AGPs in all samples was achieved with the use of the LM2 antibody, which may indicate that modification of P4H3 activity does not significantly affect the presence of β-linked GlcA because this epitope was present at a very high level at every stage of the fruit ripening process. Despite this, the silencing of the *SlP4H3* gene in the RNAi#7 line resulted in a 5-20% reduction in the intensity of the dots, depending on the stage of ripening. The most effective demonstration of the progression of the ripening process was observed in the case of JIM13 in WT. A progressive decrease in the amount of trisaccharide was found, which was also regarded as a pattern for the subsequent comparative analyses. Furthermore, an increase in the amount of the JIM13 epitope in the fruit tissue of the OEX#1 and OEX#2 lines was found. This content was 20-30% higher than in WT, even at the last stages of ripening. Also, the analysis of LM14 confirmed that more AGP epitopes were present in the transgenic lines with stimulated P4H3 activity, and the *SlP4H3* overexpression resulted in an increase in the number of LM14 epitopes. During the analysis of the LM1 epitopes, no significant differences in dot intensity were identified between the transgenic lines and WT and between the particular stages of ripening. It was also observed that the LM1 epitope was the least abundant and underwent the fastest degradation process, compared to the other epitopes.

#### 3.2.2. Molecular mass characterisation using Western blotting

The Western blotting analysis also revealed the presence of all examined epitopes of AGPs, which were characterized by different molecular weights depending on the ripening stage (Figure 3). In the WT and transgenic lines, AGPs with high molecular weight (between 120 and 60 kDa) were observed as smeared bands, which are characteristic for separation of AGPs in the electric field during electrophoresis. In the analysis of Western blot membranes, special attention was paid to the bands representing low molecular weight AGPs (∼30 kDa), which are markers of the finalisation of the ripening process in tomato fruit. The use of all the examined antibodies revealed the absence or trace quantities of low molecular weight epitopes. Interestingly, the aforementioned specific bands were not present in all the transgenic lines.

**Figure 3.**
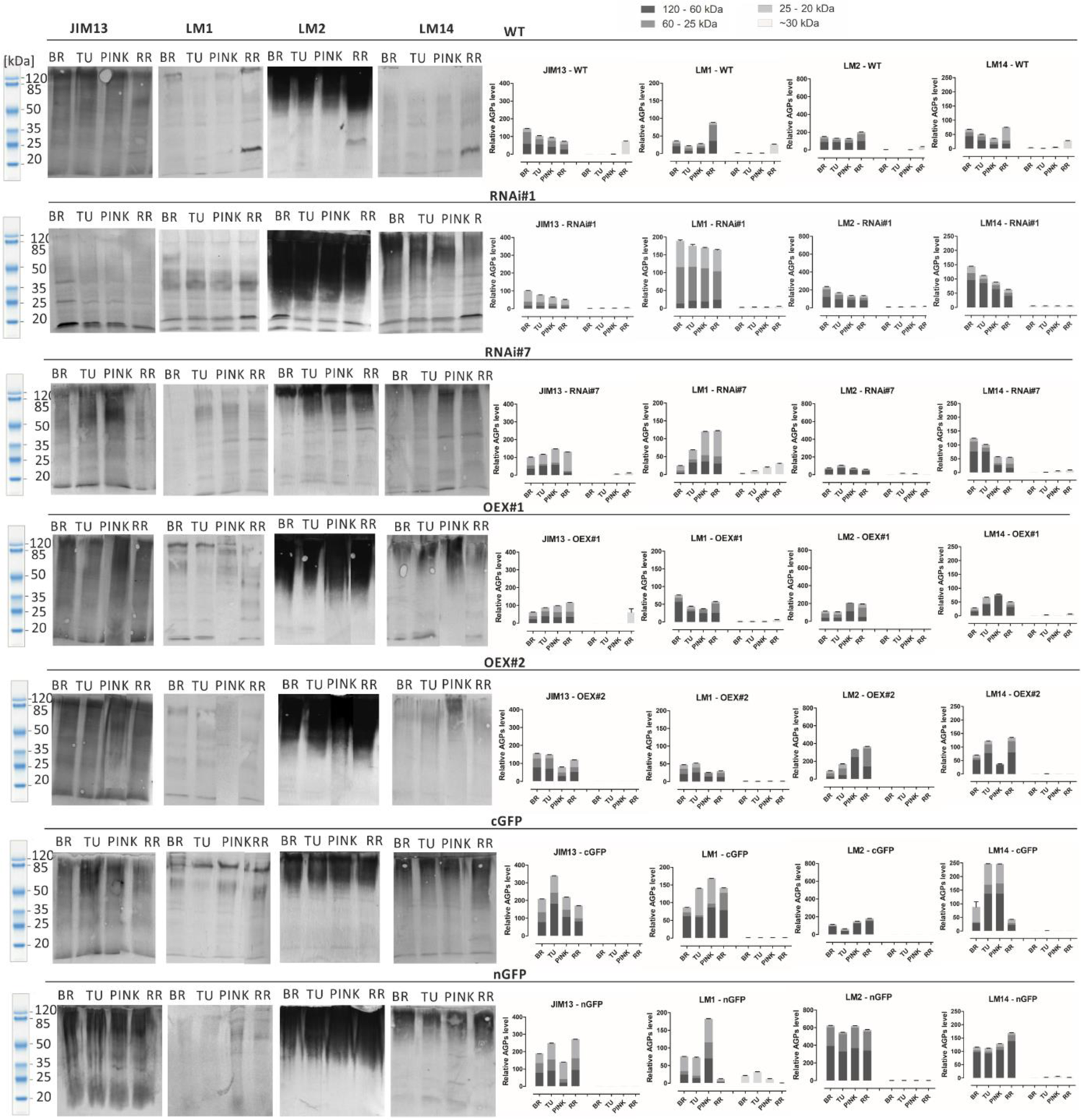
Western Blotting with quantitative analysis. Characterisation of molecular mass of AGPs from fruit at different ripening stages in wild-type plants (WT) and transgenic lines: RNAi#1, RNAi#7, OEX#1, OEX#2, cGFP, and nGFP. Specific bands around 30 kDa are marked in yellow. Abbreviations: BR – Breaker stage, TU – Turning stage, PINK – Pink stage, RR – Red Ripe stage

The next observation was that the different epitopes with various molecular weights had impaired participation in the composition of the examined lines, in comparison to WT. Both the JIM13 and LM2 epitopes were the most abundant in the WT samples and in the transgenic lines, with the highest expression level in the fruit tissue of the cGFP (JIM13 epitope) and nGFP lines (LM2 epitope). The highest LM2 expression was recorded in the nGFP transgenic line, compared to WT and the other lines. In turn, the lowest expression of the LM2 and JIM13 epitopes was observed in the fruit tissue of the RNAi#1 and RNAi#7 lines. All the transgenic lines differed from WT in the abundance of LM14 epitopes with different molecular weights. AGPs extracted from WT exhibited low expression of the LM14 epitope, while all the transgenic lines showed a significant increase in its level. The weakest participation in the sample composition in both the WT and transgenic lines was noted in the case of LM1. Only several thick bands with low molecular weight in WT (∼40kDa) and with high molecular weight (∼120 kDa) in the OEX#1 and cGFP lines were observed. The most significant result of the quantitative Western Blotting analysis showed that AGPs extracted from the transgenic lines were characterised by a more variable structure in comparison to AGPs extracted from WT. Generally, in the same ripening stages, a higher amount of AGPs with diverse molecular masses were noted in the transgenic lines than in WT.

#### 3.2.3. Selective glycome profiling of AGPs using an enzyme-linked immunosorbent assay

The results presented above identified molecular differences between samples representing the various transgenic lines, thus the ELISA test was performed to confirm the presence of the AGP epitopes and to achieve better visualisation of differences between the particular stages of fruit ripening in each line (Figure 4).

**Figure 4.**
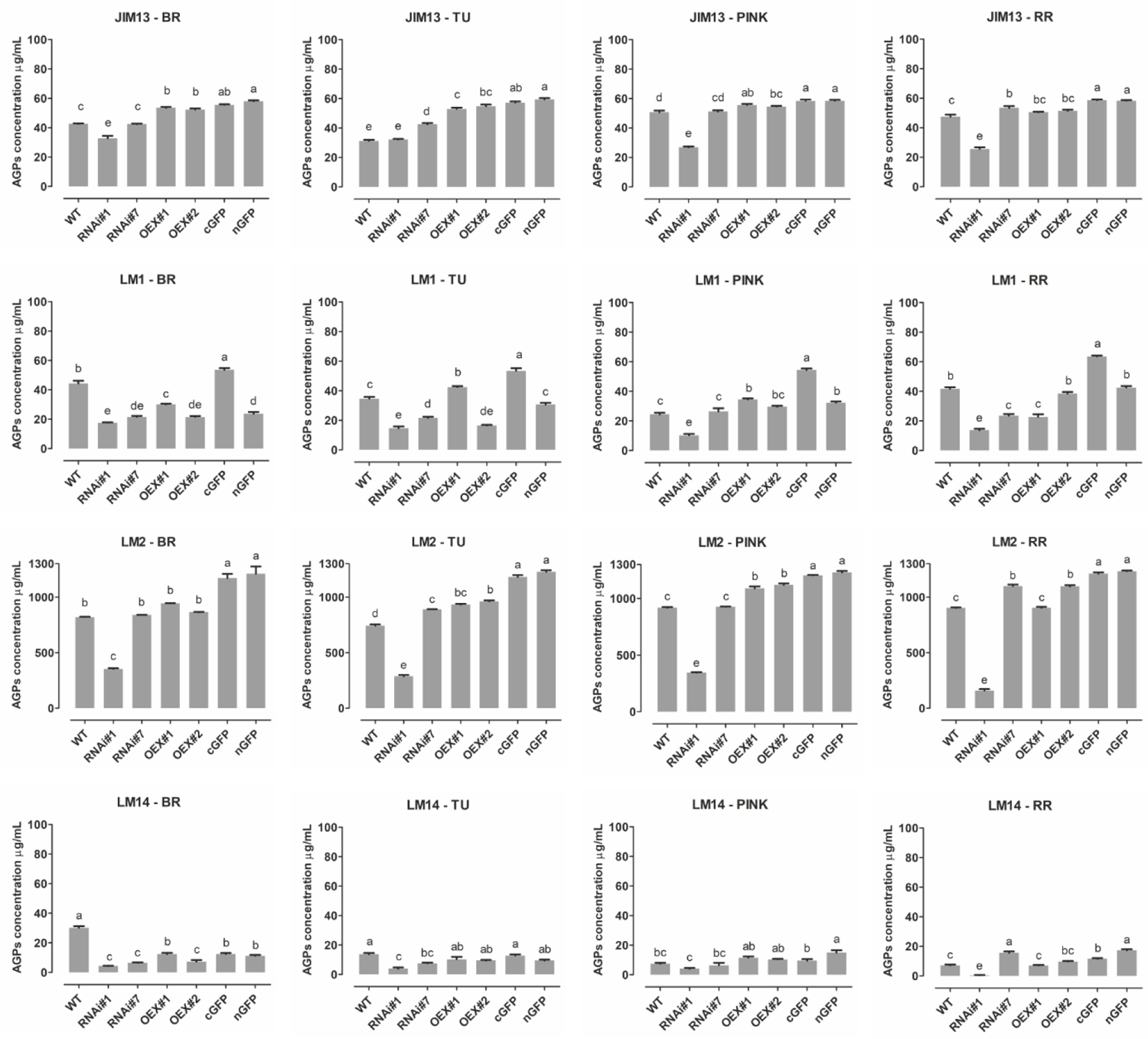
ELISA of AGPs extracted from fruit at different ripening stages from wild-type plants (WT) and transgenic lines: RNAi#1, RNAi#7, OEX#1, OEX#2, cGFP, and nGFP. Different letters indicate significant differences among the ripening stages (according to ANOVA with p<0.05). Abbreviations: BR – Breaker stage, TU – Turning stage, PINK – Pink stage, RR – Red Ripe stage

The results shown in Figure 4 indicate the lowest differences in the amount of the JIM13 epitope in the fruit of transgenic lines during the ripening process. The amount of the JIM13 epitope in the fruit tissue of the RNAi#7 line was not significantly different from that detected in WT. Besides, significant reduction of JIM13 epitope concentration in the case of RNAi#1 line. The highest concentration of the JIM13 epitope was obtained for the overexpression lines (around 55 µg/mL), and this dependence persisted to the last stages of ripening. Interesting results were obtained with the use of the LM2 antibody, as the concentration of the epitopes was significantly higher than in the analyses using the other antibodies. Only in the fruit of RNAi#1 line, the content was much lower than in other samples. Using the LM2 antibody, it was shown that the *SlP4H3* gene silencing had a significant effect on the concentration of this epitope. The AGP concentration was from approx. 700 µg/mL in WT to 1200 µg/mL for AGPs in the GFP lines, indicating that β-linked GlcA epitopes were the most abundant in the fruit AGPs. Also, the data showed that this epitope was predominant in all stages of the ongoing ripening process. In the case of overexpression of P4H3, the LM2 epitope was present in higher concentrations in comparison to native AGPs (around 20% higher for the fruit tissue of the OEX lines and around 40% for the fruit tissue of the GFP lines). The lowest absorbance value was observed after using the LM14 antibody, which correlated with the results of the dot blot analysis and Western blotting. The concentration of the LM14 epitopes had the lowest value, i.e. around 20 µg/mL in all the ripening stages and in all the tested samples. In the analyses with the LM1 antibody, significant increases in epitope concentrations were identified in the fruit tissue of the cGFP line at all the ripening stages. In turn, the fruit tissue of the RNAi#1 and RNAi#7 lines were characterised by lower LM1 epitope concentrations in comparison to WT. Also, the greatest fluctuations in the content of the LM1 epitopes were observed along with the progression of the ripening process.

### 3.3. Biochemical/ structural differences

#### 3.3.1. Estimation of AGP amount in fruit – extraction of AGPs using β-glucosyl Yariv Reagent

For biochemical analyses, AGPs were isolated using β-glucosyl Yariv Reagent. After extraction, AGPs were lyophilised and weighed. Figure 5a shows the AGP content in tomato fruit at the BR and RR stages (mg of AGPs per g of fresh tissue). This procedure made it possible to confirm that the AGP amount decreased with the ongoing ripening process both in the WT and transgenic lines. The biochemical assay using β-GlcY determined the AGP content in the tomato fruit at the BR stage: 0.38 mg/g of WT, 0.31 mg/g of RNAi#7, 4.28 mg/g of OEX#1, and 0.59 mg/g of OEX#2 and at the RR stage: 0.28 mg/g of WT, 0.29 mg/g of RNAi#7, 0.75 mg/g of OEX#1, and 0.37 mg/g of OEX#2. It revealed the most intensive degradation process in the fruit tissue of the OEX#2 line, as around 82.3% of AGPs were degraded during the progression of the ripening process. In comparison, around 26.5% in WT, 7.48% in RNAi#7, and 37.5% AGPs in OEX#1 were degraded.

**Figure 5.**
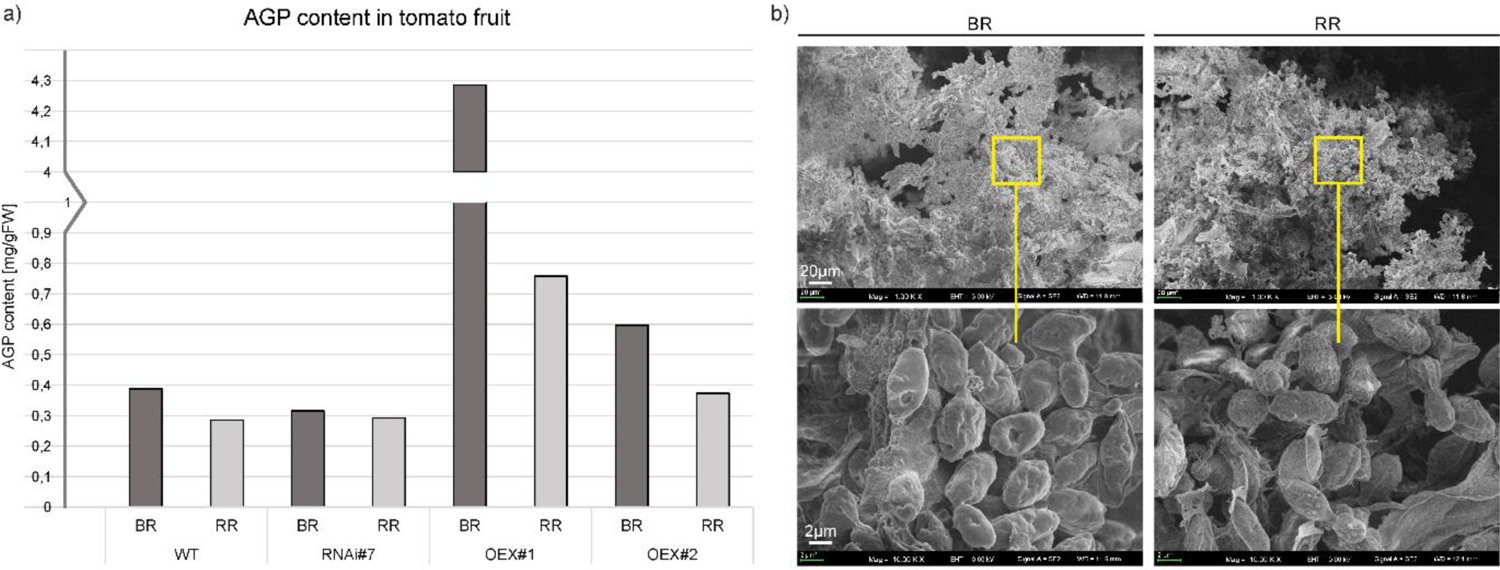
AGP content in tomato fruit at different ripening stages (a). Morphological imaging using SEM with visible AGPs and high aggregation ability (b). Abbreviations: BR – Breaker stage, RR – Red Ripe stage

#### 3.3.2. Morphological characterisation of AGPs – SEM

Figure 5b shows microphotographs of AGPs extracted from fruit at the initiation (BR) and finalisation (RR) of the ripening process. Despite the detailed observations of morphological features performed using SEM imaging, no differences between AGPs extracted from the BR and RR stages were noted. Interestingly, a high aggregation ability was visible. Thus, AGPs were imaged as compact oval aggregates composed of numerous smaller granules. The approximate diameter of the aggregates of AGPs was 1–3 µm in all the examined samples, regardless of the ripening stage.

#### 3.3.3. Structural characterisation of AGPs – FTIR

The FTIR spectra of AGPs isolated from the tomato WT and transgenic lines can be divided into several regions (Figure 6-I). The 4000–2750 region is characteristic for O-H vibrations typical for all substances and CH_2_ and CH_3_ stretching vibrations typical for lipids, proteins, and carbohydrates. The region from 1800 – 1400 is typical for bands assigned to C=O carbonyl stretching vibrations (1740 cm^-1^), carboxyl ion symmetric and asymmetric vibrations (1620 and 1416 cm^-1^, respectively), and characteristic for uronic acids. It is also typical for bands characteristic for proteins and denoted as C=O stretching in Amide I (1630 cm^-1^), as N-H bending in Amide II (1547 cm^-1^), and C-N stretching vibration in proteins ( 1̴ 450 cm^-1^) (Zhou et al., 2009; Szymanska-Chargot and Zdunek, 2013; Yamassaki et al., 2018). The region below 1400 cm^-1^ is called the fingerprint region and contains bands characteristic mostly for carbohydrates. The band at 1367 cm^-1^ is assigned to C-H vibrations and CH_2_ bending, 1331 cm^-1^ – to bending of O-H groups in the pyranose ring of pectins, 1316 cm^-1^ – to CH_2_ symmetric bending or CH_2_ rocking vibration, 1230 cm^-1^ – to bending of O-H groups in the pyranose ring of pectins and proteins, 1147 cm^-1^ – to ring vibrations (C-OH) overlapped with stretching vibrations of side groups and glycosidic bond vibrations (C-O-C), 1000–1200 cm^-1^ – to skeletal vibrations of the ring or the glycoside bond, 1019 cm^-1^ – to C-O stretching and C-C stretching (Boulet et al., 2007b; Szymanska-Chargot and Zdunek, 2013; Szymanska-Chargot et al., 2015; Chylińska et al., 2016). The bands at approx. 896, 835, and 776 cm^-1^ are characteristic for β-glycosidic linkages (hemicelluloses and cellulose), α-glycosidic linkages (uronic acids), 1–4 linkage of galactose, and 1–6 linkage of mannose (Bashir and Haripriya, 2016). The bands below 700 cm^−1^ are attributed to the skeletal vibrations of pyranose rings (Szymanska-Chargot et al., 2015; Bashir and Haripriya, 2016).

**Figure 6.**
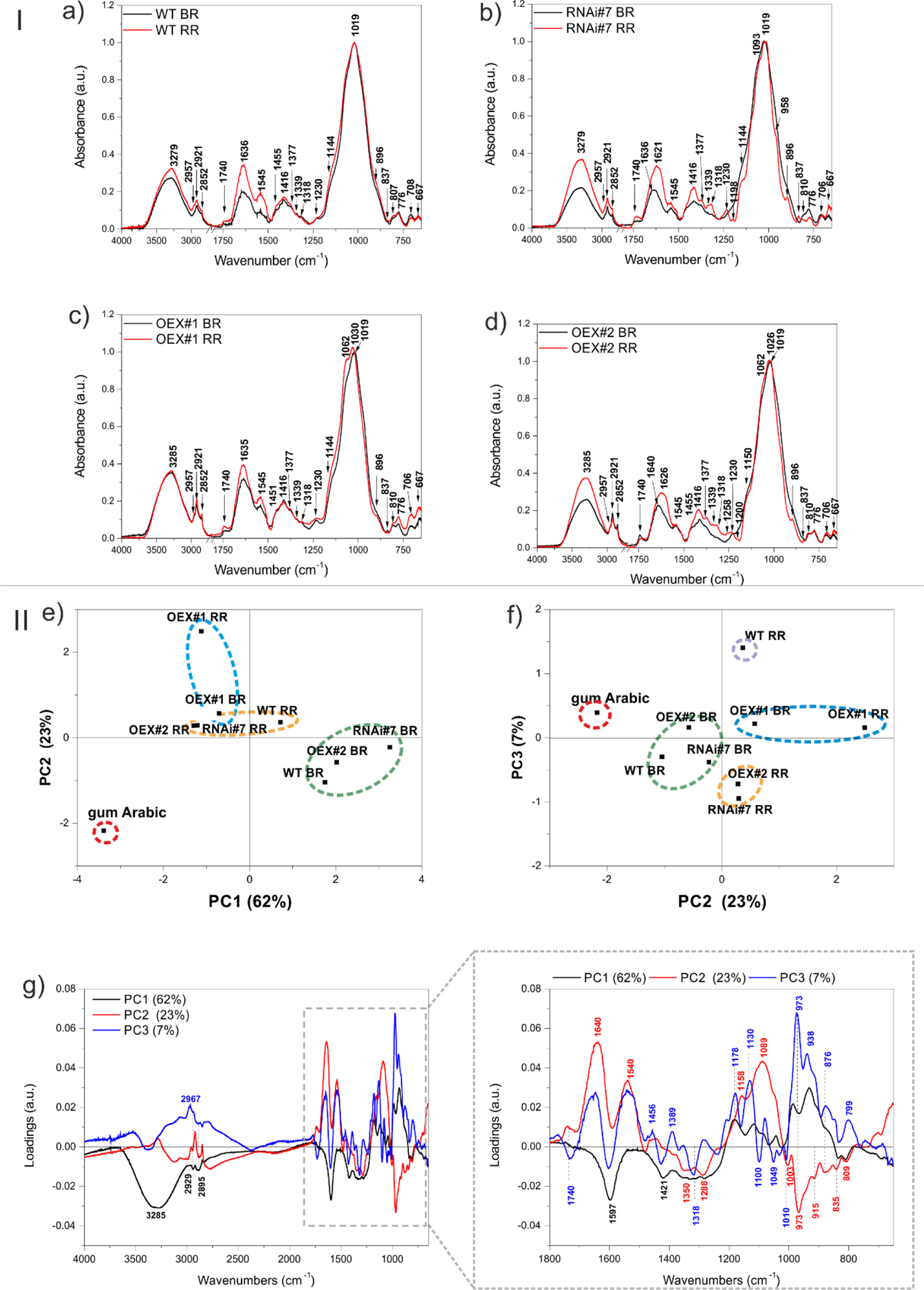
I. FTIR spectra in the range of 4000 – 650 cm^-1^ of AGP isolates obtained from tomato fruit at different ripening stages in wild-type plants (WT) (a) and transgenic lines: RNAi#7 (b), OEX#1 (c), and OEX#2 (d). The 2750 – 1800 cm^-1^ region was cut out as it did not show any spectral features. The most characteristic wavenumbers are highlighted in the spectra. **II.** PCA of the FTIR spectra of AGP isolates from tomato fruit at different ripening stages in wild-type plants (WT) and transgenic lines: RNAi#7, OEX#1, and OEX#2. Gum Arabic was added to the analysis as the AGP standard. Scatter of PC1xPC2 (e) and PC2xPC3 (f) scores are shown together with the loading profile of components PC1, PC2, and PC3 (g). The most characteristic wavenumbers are highlighted in the loadings; different colours (black, blue, and red) denote the different PC (PC1, PC2, and PC3) loadings, respectively (some of the wavenumbers are characteristic for more than one PC loading). Abbreviations: BR – Breaker stage, RR – Red Ripe stage

In the case of the AGPs isolated from the BR and RR stages of WT, the FTIR spectra did not show substantial differences (Figure 6a). Generally, in addition to arabinogalactans (β-glycosidic linkage vibrations at 776 cm^-1^) and proteins (amide bands at 1630 cm^-1^, 1547 cm^-1^, and 1450 cm^-1^), the isolates also contained uronic acids and pectins (bands at 1367 cm^-1^, 1331 cm^-1^, 1316 cm^-1^, 1230 cm^-1^, and especially 835 cm^-1^ characteristic for α-glycosidic linkages).

In the case of the RNAi#7 line, the differences in the FTIR spectra between the BR and RR stages were apparently more pronounced (Figure 6b). The spectrum for the BR stage of the RNAi#7 line was very similar to the BR stage of WT with a difference that the band at 1740 cm^-1^ for the mutant was more intense than for WT, which may evidence the presence of more esterified pectins. This band was even more intense in the spectrum for the RR stage of the RNAi#7 line. The spectrum for the RR stage of RNAi#7 line bands at 1377, 1339, 1318, 1230, 1093, 958, and 837 cm^-1^ characteristic for pectic polysaccharides, especially rhamnogalacturonans type I and II (Boulet et al., 2007), and 896 cm^-1^ characteristic for most of the hemicellulosic polysaccharides increased, while those characteristic for pure arabinogalactan glycoprotein (Arabic gum spectrum, SM Figure 6I) at 810 and 776 cm^-1^ decreased (Szymanska-Chargot and Zdunek, 2013).

Less clear differences between spectra were obtained for the BR and RR stages in the OEX#1 line (Figure 6c). The spectra were characterized by relatively high intensity of the band at 1740 cm^-1^ assigned to esterified pectins. In turn, the bands at 1062 and 1030 cm^-1^ that appeared in the spectrum for the RR stage of the OEX#1 line can be assigned to the vibrations of pyranose in gum Arabic and arabinogalactan polysaccharide, respectively (Wu et al., 2020). Moreover, the band at 837 cm^-1^ characteristic for pectic polysaccharides diminished and bands at 896, 810, and 776 cm^-1^ characteristic for arabinogalactans were more visible in the spectrum for the RR stage of the OEX#1 line.

Interestingly, the band at 1740 cm^-1^ declines and the band at 1640 cm^-1^ is shifted to 1626 cm^-1^ in the spectrum for the RR stage of the OEX#2 line, compared with the BR stage of the OEX#2 line (Figure 6d). This may evidence demethylation of pectic polysaccharides present in the AGP isolate from the OEX#2 line. Additionally, as in the case of the RR stage of the RNAi#7 line, the bands characteristic for pectic polysaccharides, especially rhamnogalacturonans type I and II, also appeared in the RR stage of the OEX#2 line. However, the AGP isolate from the RR stage of the OEX#2 line was also rich in arabinogalactan polysaccharide and protein, as the bands visible at 1150, 1062, 1026, 896, 810, and 776 cm^-1^ were intense.

The PCA analysis of the obtained FTIR spectra was performed for better comparison and highlighting of the samples (Figure 6-II). Figures 6e and 6f present the scatter plots of the scores of the first three principal components PC1, PC2, and PC3, which together explain 92% of variability. The scores are scattered along all axes and the grouping effect of scores can be observed. The gum Arabic is separated from the other samples, which is clear evidence that the spectra of the obtained isolates differ from the AGP standard (Figure 6e, 6f). The samples from the BR stages of WT, RNAi#7, and OEX#2 and samples from the RR stages of WT, RNAi#7, and OEX#2 form two separate clusters in the PC1xPC2 score plot, which proves that there are spectral similarities between the samples, while the samples from the BR and RR stages of the OEX#1 line form a separate cluster (Figure 6e). In the PC2xPC3score plot, the same samples groups were obtained; however the RR stage of WT form a new separate group (Figure 6f).

Loadings that influence the differentiation of the scores are presented in Figure 6g. The wavenumbers of PC1 negative loadings that mostly influenced the sample scores lying on the negative side of PC1 (gum Arabic, the BR and RR stages of the OEX#1 line, the RR stages of the OEX#2 and RNAi#7 lines) were 3285, 2929, 2895, 1597, 1421, 1350, 1018, 1288, 835, and 810 cm^-1^, which are mostly characteristic for esterified pectins and for AGP protein (gum Arabic spectrum), while the positive influence was associated with the following wavenumbers: 1181, 1119, 1042, 977, and 938 cm^-1^, mostly characteristic for gum Arabic (SM Figure 6-I). On the other hand, the gum Arabic was located on the negative side of PC2, on which wavenumbers 1350, 1288, 973, 835, and 809 cm^-1^, also characteristic for pure AGP from gum Arabic, had an influence. The other samples lying on the negative side of PC2 (but closer to PC2=0) were the BR stages of WT, OEX#2, and RNAi#7 lines. The wavenumbers having a positive influence on the scores of samples scattered on the positive side of PC2 were 1740, 1640, 1540, 1158, and 1089 cm^-1^. The negative influence on the scores scattered along PC3 (the RR stages of the OEX#2 and RNAi#7 lines) had wavenumbers 1740, 1597, 1421, 1318, 1100, 1049, and 1010 cm^-1^ – characteristic mostly for pectic polysaccharides. In turn, wavenumbers 1178, 1130, 973, 938, 876, and 799 cm^-1^, characteristic for arabinogalactans and for some phospholipids, had a positive influence.

### 3.4. Differences in AGP arrangement in the cell wall assembly – analyses *in situ*

#### 3.4.1. Localisation of AGPs at the cellular level (CLSM)

*In planta* studies were performed using the immunofluorescence technique to identify presumed changes in the spatio-temporal pattern of the AGP distribution in the fruit tissue of the examined lines. Figures 7, 8, and 9 show microscopic investigations of the AGP arrangement in the tissue of WT, OEX#1, and OEX#2; RNAi#1, and RNAi#7; cGFP and nGFP lines, respectively. Our decision to focus on the two distant stages of ripening (BR and RR) was based on the fact that the differences did not appear between particular successive stages.

**Figure 7.**
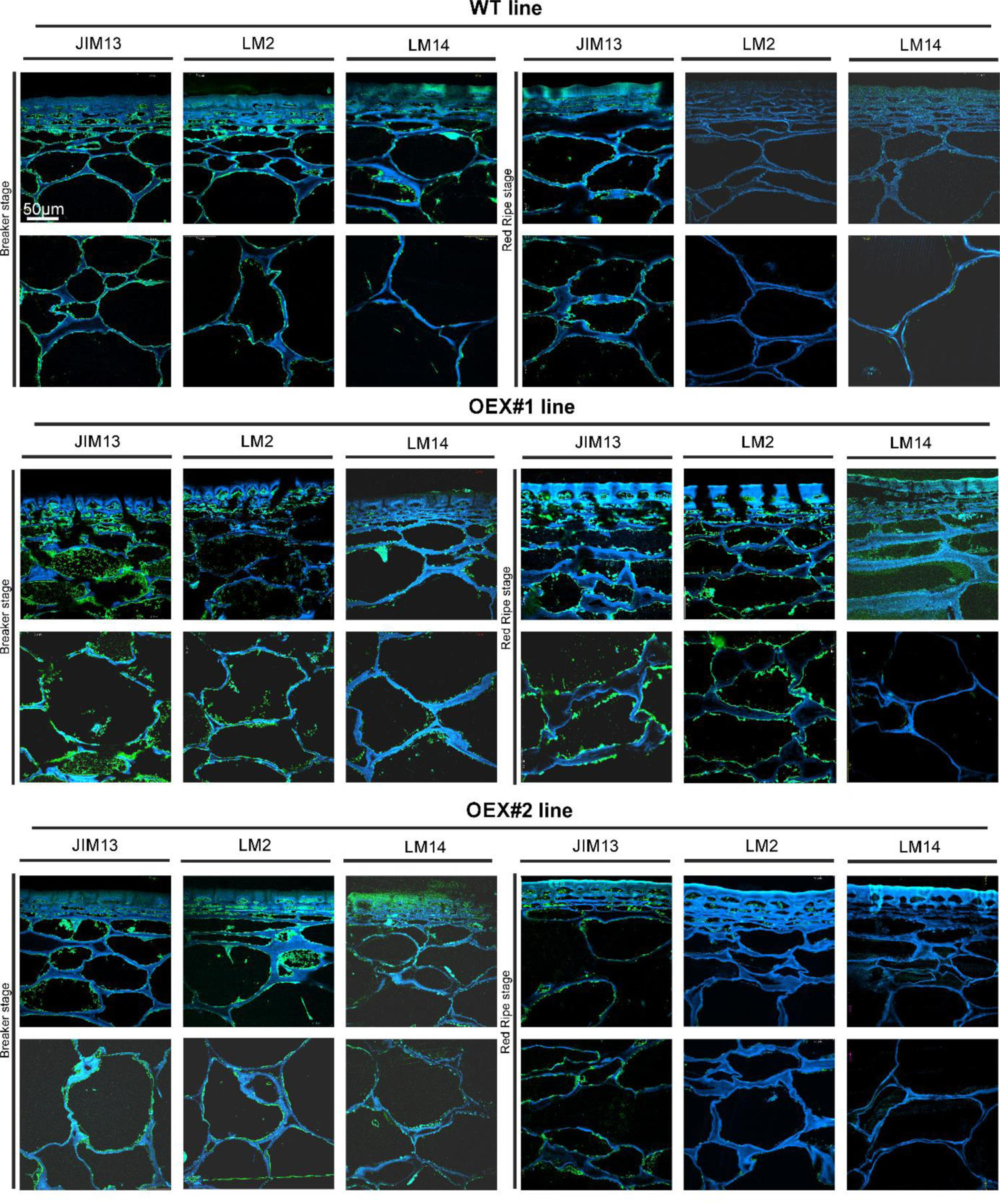
Spatio-temporal localisation of AGPs in tomato fruit tissue at different ripening stages in wild-type plants (WT) and transgenic lines OEX#1 and OEX#2. Immunofluorescence method using JIM13, LM2, and LM14 antibodies.

**Figure 8.**
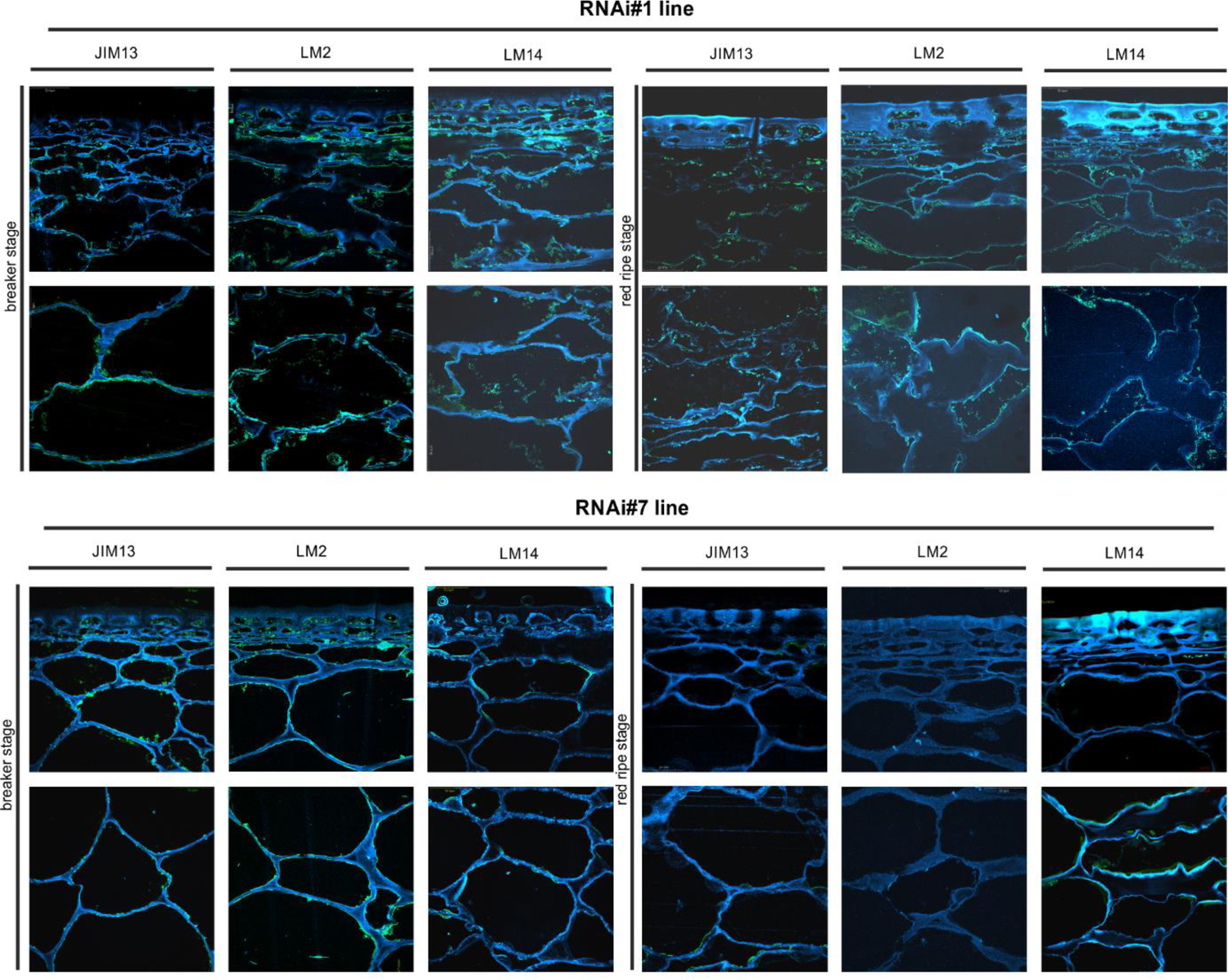
Spatio-temporal localisation of AGPs in tomato fruit tissue at different ripening stages in transgenic lines RNAi#1 and RNAi#7. Immunofluorescence method using JIM13, LM2, and LM14 antibodies.

**Figure 9.**
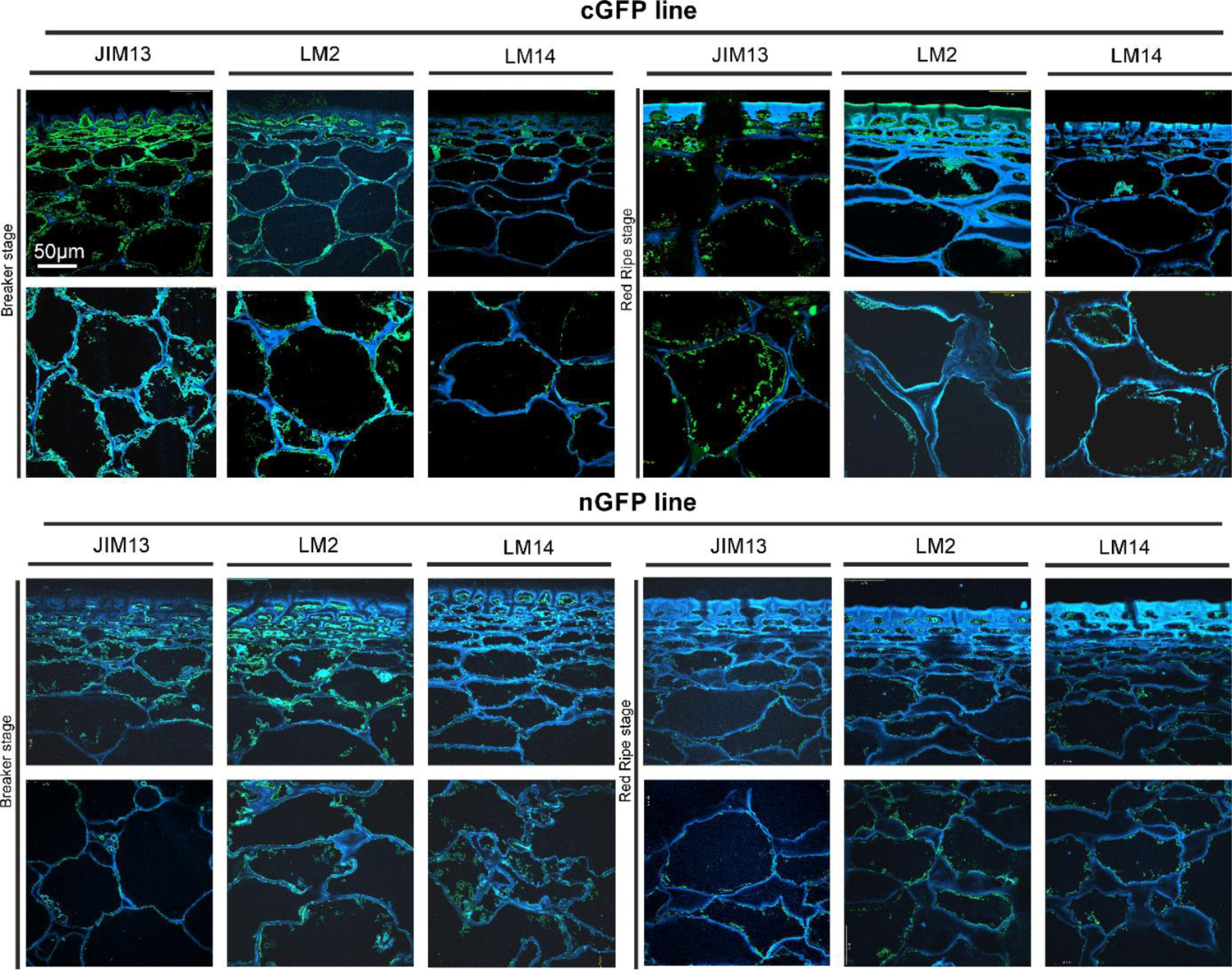
Spatio-temporal localisation of AGPs in tomato fruit tissue at different ripening stages in transgenic lines cGFP and nGFP. Immunofluorescence method using JIM13, LM2, and LM14 antibodies.

The microscopic observations demonstrated the anatomic changes in fruit tissues and cell walls at the cellular level. In all the fruit lines, modifications of cell walls occurred with the progression of the ripening process and were visualised as degraded cellular compartments adjacent to swollen cell walls. These modifications were correlated with typical processes of decay in fruit during the ongoing ripening process. The characteristic pattern of AGP distribution as a marker of ripening progress, described in detail in our previous paper, was observed in the fruit tissue of WT at the BR stage (Kutyrieva-Nowak et al., 2023a). The specific pattern of arrangement of AGPs was labelled as a line at the border between the cell wall and the plasma membrane. With the subsequent stages of ripening, the localisation of AGPs changed and the immunofluorescence intensity was lower. For WT, this correlation persisted for the samples labelled with all the antibodies.

The interpretation of microscopic results consisted in finding differences between the analysed lines after the reactions with three antibodies against AGPs. In the initial ripening stages, the fluorescence intensity in the OEX#1 fruit tissue was higher than in WT. The increased number of labelled AGP epitopes was confirmed using all antibodies. The fruit tissue of the OEX#2 line also showed an increase in the number of labelled AGP epitopes compared to WT, although the number was lower than in the fruit tissue of the OEX#1 line. The increase in the amount of AGP epitopes (mainly JIM13 and LM2) disturbed the typical pattern of localisation of AGPs, and the epitopes were visible in the whole area of the cell wall-plasma membrane continuum. Similarly, the striking typical anatomical changes in the tissue during the ripening process coincide with a decrease in the occurrence of AGP epitopes. However, the immunofluorescence signal is clearly higher compared to WT.

In samples with the silenced expression of the *SlP4H3* gene, intrusive alterations in the tissue morphology were imaged as a result of ripening progress. Significant changes in the macrostructure were found in the fruit tissue of the RNAi#1 and RNAi#7 lines. At first, compared to all lines, the tissue in the mutants mentioned above was looser even at the BR stage. In addition, at the RR stage, the tissue was disordered, the cell wall was less integrated, unevenly thickened, more undulating, and swelled. The rearrangement of epitopes was correlated with cellular dissociation. In the fruit tissue of the RNAi# lines, a decrease in all AGP epitopes was observed at both the BR and RR stages.

In the fruit tissue of the cGFP line, substantially increased fluorescence intensity was observed after the reaction performed using JIM13, LM2, and LM14, compared to WT. Moreover, AGP epitopes were present at the last stage of ripening. In the fruit tissue of the nGFP line, loosening of the cell wall was observed from the first ripening stage. With the progression of the ripening process, the cell wall became even more unevenly undulating and thickened. The number of AGP epitopes decreased, which distinguished this line from the fruit tissue of the OEX and cGFP lines.

#### 3.4.2. Localisation of AGPs at subcellular level (TEM)

The immunogold labelling analysis and TEM imaging (40 kx magnification) allowed localisation of AGPs in the individual compartments of the fruit cell at the subcellular level. The analysis was carried out using the JIM13 antibody. Figure 10 shows analyses of the correlation between the localisation of AGPs in the cellular compartments and the stages of the ripening process. JIM13 epitopes, visible as dark points marked with red circles (Figure 10a), were observed in the entire cell, and labelling mainly occurred in the cell wall-plasma membrane continuum and the cytoplasm. For consistency, we analysed two cell compartments (cell wall-plasma membrane and cytoplasm) in all cases. Since there were no significant differences between all the subsequent stages of the ripening process noted by CLSM imaging at the cellular level, the analysis at the subcellular level was carried out in two distant stages of ripening (BR and RR). Generally, AGP epitopes were detected less frequently in the RR stage of ripening than in the BR stage. In almost every case, the cell wall-plasma membrane continuum exhibited the largest amounts of AGP epitopes, with the exception of the RR stage in the fruit tissue of the cGFP and nGFP lines.

**Figure 10.**
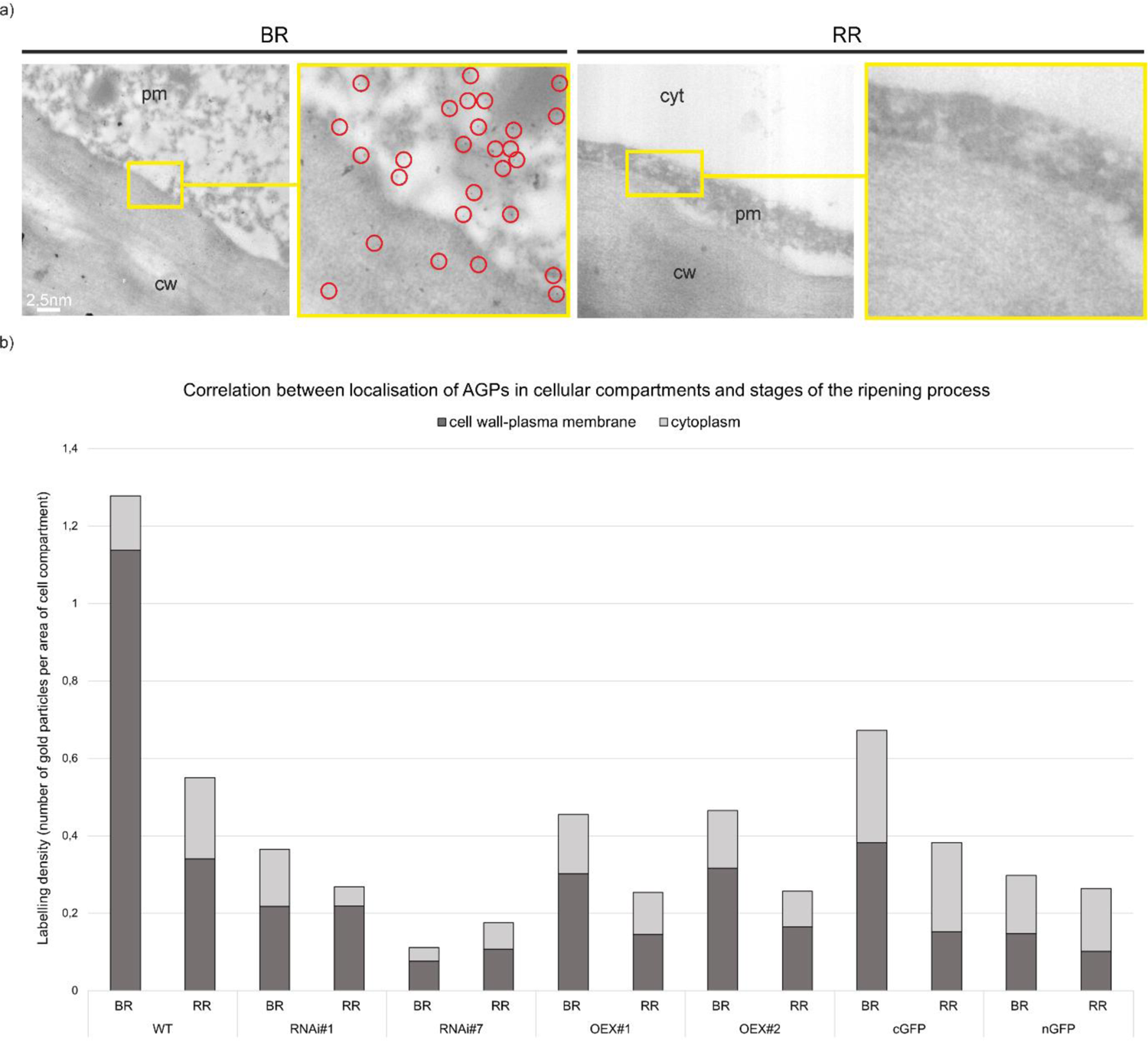
Ultrastructural TEM imaging of immunogold labelling of AGPs in tomato fruit at different ripening stages. Gold particles marked with red circles (a). Quantitative data derived from analyses of the density of gold particles (b). Abbreviations: cw – cell wall, pm – plasma membrane, cyt – cytoplasm, BR – Breaker stage, RR – Red Ripe stage

The quantitative analysis of immunogold labelling confirmed the specific spatio-temporal pattern of distribution of AGPs and indicated changes in the amount of AGPs in the cell wall and cytoplasm in the transgenic lines during the ripening process (Figure 10b). Once again, the most characteristic distribution of AGPs was obtained in the WT fruit tissue. At the BR stage, the AGP epitopes were more organized and localised mainly in the cell wall-plasma membrane continuum. Only 10% of gold particles were localised in the cytoplasm. However, during the ripening process, the number of gold particles increased in the cytoplasm and accounted for nearly 40% at the RR stage. This may evidence the process of degradation of the cell wall and release of residues of cell wall compartments into the cytoplasm.

The AGP distribution pattern was disrupted in the fruit of the transgenic lines, where AGPs were dispersed in the whole area of the extracellular matrix. It was shown that modifications of *SlP4H3* gene activity caused changes in the number of gold particles in particular cell parts. Moreover, different proportions of gold particles in the cell wall-plasma membrane and in the cytoplasm were found in the examined samples, compared to WT. In lines with the overexpression of the *SlP4H3* gene, an increased amount of AGPs in the cytoplasm was found. Furthermore, the subcellular localisation of the AGP epitope in the fruit tissue of the cGFP and nGFP lines at the BR stage showed the presence of half of the amount of gold particles in the continuum cell wall-plasma membrane (0.38 and 0.14 particles per µm^2^, respectively), and the rest was distributed in the cytoplasm (0.29 and 0.15 particles per µm^2^, respectively). On the other hand, a significant reduction in the number of gold particles in the cell wall-plasma membrane (almost 80%) was observed in the lines with silenced *SlP4H3* gene expression (RNAi#1 and RNAi#7) at the BR stage. Similarly, at the RR stage, the AGP assembly disorder was assessed, and a higher amount of gold particles was noted in the cytoplasm than in the cell wall-plasma membrane.

## 5. DISCUSSION

The ripening process is the subject of extensive research, but the role of AGPs in this process is still poorly known (Seymour et al., 2013). In our research, the genetic modification in the AGP biosynthesis process by changes in the activity of the P4H enzyme allowed identification of AGP alterations during the ripening process. The results obtained allow concluding that changes in the expression of the *SlP4H3* gene affect AGPs and this, in turn, generally affects the cell wall structure. Disruption in hydroxylation and subsequent AGP glycosylation processes most probably induce a sequence of modifications and degradation of the proper assembly in the arrangement of cell walls. In the current work, we did the work step by step to investigate how changes in the process of forming the protein moiety may influence the subsequent phase of the AGP molecule synthesis pathway, such as attachment of carbohydrate chains. The analyses performed using molecular, structural, and microscopic tools in *ex situ* and then *in situ* studies revealed crucial changes in AGP features in the fruit of the transgenic lines in comparison to the wild-type plants. Based on the results obtained, a meticulous pattern of consecutive events in fruit cells that could be associated with P4H3 was developed.

The development of the overexpressing and silenced P4H3 tomato lines facilitated the determination of alterations in the AGP carbohydrate design that induced changes in their localisation in the cell wall, allowing identification of AGP structural functions in the fruit tissue. So far, it has been demonstrated that hydroxyproline-rich glycoproteins are involved in fruit ripening (Moore et al., 2014). One of the major findings of a study conducted by Moore and coworkers on the anti-extensin LM1 antibody showed an increasing amount of extensin epitopes in ripe grapes as a response to the ripening process (Moore et al., 2014). These observations were consistent with the results of the analysis carried out by Nunan and coworkers that protein components rich in arginine and hydroxyproline also accumulate in grapes at the last stages of ripening (Nunan et al., 1997, 1998). Similarly, our previous studies on apple and tomato fruit have shown that the amount of AGPs during the fruit ripening process changed with a significant concentration increase at the particular stages of the ripening programme (Leszczuk et al., 2020c; Kutyrieva-Nowak et al., 2023a). The analysis carried out with the use of β-glucosyl Yariv Reagent, whose reactivity is one of the most important criteria in the identification of AGPs (Seifert and Roberts, 2007; Kitazawa et al., 2013; Tsumuraya et al., 2019), also confirmed the differences in the AGP content between the specific stages of ripening. In the study conducted by Peng and coworkers, the β-glucosyl Yariv Reagent gave a strong positive reaction for AGPs fraction from ripe fruit of *Lycium ruthenicum* Murr, which is a water-soluble AGP (Peng et al., 2012). The confirmation was found also in the present work, as the content of AGPs decreased with the ongoing ripening process both in WT and in all the examined transgenic lines. Interestingly, an increased amount of AGPs in the overexpression lines was observed, indicating the role of P4Hs in the AGP biosynthetic pathway. However, the degradation process in these lines progressed more intensively than in WT and lines with silenced gene expression.

The next research step involved comprehensive structural analyses on extracted AGPs aimed at revising changes that may have appeared as a result of the P4H3 modifications. The FTIR technique used for this purpose provided information about carbohydrate chains in AGPs during the fruit ripening process. Previous single FTIR reports of monosaccharide compositional analysis of AGPs isolated from *Lycium ruthenicum* Murr fruit revealed that arabinose and galactose represented by absorbance regions at 1060-1040, 975 cm^−1^, and 945 cm^−1^, respectively, were the major AGP sugars (Coimbra et al., 1998). The results obtained in the current work also confirm the occurrence of these carbohydrates in the isolated AGPs. The FTIR spectra showed that the AGP isolates from tomato fruit contained pure arabinogalactan proteins and a trace of pectic polysaccharides (uronic acids, rhamnogalacturonan) and hemicelluloses. The PCA analysis in the current work identified three PC groups with different polysaccharide compositions. The result obtained for the analysis of AGPs from the OEX#1 line seems interesting, as samples collected at different ripening stages formed a common group, compared to the two separate groups of the BR and RR stages. Besides, the PC2xPC3 analysis identified a disruption of the ‘native’ ripening pattern of the transgenic lines, as they did not form a common group with the WT RR. In both PCA analyses, samples at the BR stage from the OEX#2 and RNAi#7 lines formed one group with WT. We may conclude that the modification of *SlP4H3* significantly affects the progression of the ripening process, causing changes in the profiles of the components of AGPs.

Molecular studies were performed for verification of the differences between the tomato lines in detail. In contrast to the β-glucosyl Yariv Reagent assays, anti-AGP antibodies recognizing specific carbohydrate epitopes used in these methods constitute a differentiating factor (Gao et al., 1999). Firstly, the immunoprinting on the nitrocellulose membrane confirmed the presence of AGP epitopes in tomato extracts and highlighted that the overexpression lines were characterised by a general trend towards increased dot intensity, indicating higher content of the tested AGP epitopes. Secondly, a detailed analysis of the composition of individual epitopes in the analysed lines showed further differences indicating disturbance in the carbohydrate chains of AGPs as a result of the modifications in P4H3 activities. In the study conducted by Fragkostefanakis and coworkers on tomato pericarp tissue with a ripening process without disorders, changes in the amount of AGP epitopes and a significantly higher number of the JIM13 epitope were noted (Fragkostefanakis et al., 2012). Western blotting analyses of the JIM13 epitopes revealed a slight decrease in high molecular weight epitopes in the last ripening stages. In the case of the JIM8 antibody, a gradually decrease in the presence of AGPs was also observed after the turning stage (Fragkostefanakis et al., 2012). On the other hand, in the study carried out by Sun and coworkers, two major and three minor peak fractions were collected after biochemical separation of AGPs from tomato by RP-HPLC (Sun et al., 2004). Western blotting with the LeAGP-1 antibody verified that only one fraction contained AGPs with a molecular weight of 48 kDa (Sun et al., 2004). In the first report on the molecular mass of AGPs extracted from fruit by Fragkostefanakis and coworkers, AGP epitopes recognised by JIM8 and JIM13 epitopes in tomato pericarp tissue were in the range of 210-55 kDa and 300-45 kDa, respectively (Fragkostefanakis et al., 2012). However, in our previous papers, the molecular mass of AGPs was about 250-70 kDa from apple fruit (Leszczuk et al., 2020c) and 120-20 kDa from tomato fruit (Kutyrieva-Nowak et al., 2023a), which may indicate a variable pattern of AGPs in different fruits. In addition, in the case of the JIM13 epitope in WT at the beginning ripening stages, AGPs with 120–60 kDa and 60–25 kDa molecular weight predominated, and at the last stages contained 30–20 kDa AGPs, which were considered a marker of the finalisation of the ripening process in tomato fruits (Kutyrieva-Nowak et al., 2023a). In the current work, AGPs from the transgenic lines were characterised by an altered molecular mass in comparison to WT. The molecular differences and the absence of single low molecular weight bands in the last stages of ripening indicate an effect of the P4H3 activity disruption on AGP structural modifications. The changes observed also pointed to the disturbed process of degradation of ‘native’ carbohydrate chains with the ripening progress. Selective glycome profiling of AGPs using an enzyme-linked immunosorbent assay allows rapid analysis of carbohydrate epitopes (Pattathil et al., 2010). Data from previous papers provided information about the relative absorbance of AGP epitopes for the JIM13 antibody. The strength of ELISA signals was characterised by absorbance at 0.05-0.08 in the case of banana fruit and at 0.71-1.18 in the case of mango fruit (Rongkaumpan et al., 2019). Research on tomato fruit from wild-type plants showed an average absorbance level of 0.5-3.5, with significant changes correlated with the ongoing ripening process (Kutyrieva-Nowak et al., 2023a). The present data showed that the absorbance level dependent on the tested lines was different from that of AGPs extracted from fruit in previous studies. This mainly involved the use of JIM13 and LM2 antibodies, where we obtained absorbance levels at 2.5-4 and 2.9-4.2, respectively. The results obtained with the LM14 antibody seemed interesting as the absorbance of the transgenic lines was 0.5-0.9 compared to the absorbance of around 2.0 for WT. The glycome profiling with the LM1 antibody did not allow definite determination of the dependence of absorbance changes during the ripening process, as it was quite variable at the different stages of ripening. The changes found were related to disruptions in the concentration of specific epitopes caused by modifications in *SlP4H3* gene expression. Interestingly, the increase in the AGP concentration in the overexpression lines and the decrease in the AGP concentration in the lines with silenced P4H3 expression were confirmed by this method as well. All structural and molecular approaches used in this part of the work clearly indicate a linkage between P4H3 activity and the AGP carbohydrate moiety structure.

*Ex situ* studies showed that AGPs extracted from the examined transgenic lines were characterised by different molecular structures of carbohydrate chains, which is also connected with particular alterations linked with the stages of ripening. Our next goal was to investigate how the structural changes affected the AGP localisation in the cell wall assembly. This issue is even more interesting and necessary to analyse, as previous studies have identified the characteristic spatio-temporal pattern of AGP distribution in the cell wall-plasma membrane continuum at the cellular and subcellular levels (Leszczuk et al., 2018; Kutyrieva-Nowak et al., 2023a). For example, in a study on tomato LeAGP-1, Sun and coworkers observed AGPs from the cell wall bound with the plasma membrane via a glycosylphosphatidylinositol anchor (Sun et al., 2004). This allows AGPs to function as a linker between the plasma membrane and the cell wall (Ellis et al., 2010; Liu et al., 2015; Ma et al., 2017). Next, subcellular studies of fruit confirmed the information about the distribution of AGP epitopes at the border of the cell wall-plasma membrane continuum and proved that AGP epitopes changed their location and demonstrated a decrease in the number of AGP epitopes with the progression of the fruit senescence process, which is linked to degradative processes in the cell wall and the release of its cellular compartments (Leszczuk et al., 2018). Moreover, it is well-known that the variable presence of specific epitopes in different stages shows dynamic changes in the cell wall during the ripening processes. Research conducted by Szymańska-Chargot and coworkers provided information on changes in polysaccharide distribution during post-harvest apple storage (Szymańska-Chargot et al., 2016). Comprehensive microarray polymer profiling performed by Moore and coworkers demonstrated an increase in JIM13 and LM2 epitopes during the ripening process of grapes; the signal intensity was significantly weaker in green grapes compared to ripe grape berries (Moore et al., 2014). This was also confirmed in our previous research, where changes in the AGP distribution pattern were observed during the development and ripening processes (Leszczuk et al., 2018, 2020c; Kutyrieva-Nowak et al., 2023a). In apple fruit tissue, the amount of AGPs during the senescence process in post-harvest storage also decreased, which was accompanied by a noticeable decrease in the immunofluorescence intensity (Leszczuk et al., 2020c). The ripening-associated changes had an impact on not only the localisation and distribution of AGPs and other cell wall components but also the fruit texture, causing differences in the anatomical features in the fruit tissue (Winisdorffer et al., 2015; Leszczuk et al., 2019b). This confirms that the correct glycosylation of AGPs is essential for proper cell wall assembly and extracellular matrix functioning (Velasquez et al., 2011). Also, the analyses conducted in the present work revealed a decrease in the total number of epitopes during the ripening process with a concomitant increase in AGPs in the cytoplasm, compared to the cell wall-membrane continuum. Nevertheless, the significant disruption of the specific and well-documented AGP localisation pattern in all the examined transgenic lines must be underlined. In addition, the disorder of the proportion of AGPs attached to the cell wall and placed in other cellular compartments was also observed for all the transgenic lines. Moreover, numerous observations at the tissue level allow concluding that the overall appearance and structure of fruit tissue are also modified. The visible different localisation of AGPs related to the structural and molecular disruption caused by the change in the P4H3 activity is connected with modifications of fruit tissue morphology.

All analyses performed in the present study create complementary evidence that the effect of modifications of *SlP4H3* gene expression is supported by changes in the AGP amount, structure, and compositional properties, and thus impaired distribution of AGPs in fruit cells.

## 6. CONCLUSIONS

To sum up, the study demonstrated the disturbance caused by the modifications, i.e. changes in the AGP content, disruption of AGP molecular mass, alterations in the molecular composition, changes in the amount of AGPs bound by Yariv Reagent, structural differences indicating various degrees of ripening progress, and the disordered distribution pattern in the cell wall, plasma membrane, and cellular compartments.

Based on the obtained results, we focused on presenting the sequence of events that may take place in the fruit cells in response to changed P4H3 activity:

1. Disturbance of the ‘native’ AGP structure. The changed content of hydroxyproline residues affects the protein moiety of AGPs, which is associated with altered attachment of carbohydrate chains;
2. Disruption of the AGP molecular structure (mainly carbohydrate chains) causes changes in their localisation at the subcellular levels;
3. Changes in the spatio-temporal AGP distribution may be related to the formation of a stable network between AGPs with other cell wall components (APAP1).
4. The lack of properly formulated AGP chains affects the interlinking in the APAP1 network. In turn, the modified stability of the cell wall assembly has an influence on anatomical changes visible at the cellular level.
5. Changes in the continuity and durability of the cell wall affect the entire fruit tissue and the sequence of tissue transformation during the progression of the ripening process.

We conclude that the impaired P4H3 activity has an effect on the AGP molecular and structural features and, consequently, affects the degradation of AGPs during the ongoing ripening process. We may suppose that the ‘native’ structure of AGPs, mainly their carbohydrate moieties, is important for the strictly scheduled extracellular matrix arrangement and is essential for the correct course of the fruit ripening process. An interesting result is that the modifications caused changes in the anatomical image of the entire fruit tissue. In subsequent studies, it is necessary to focus on other components of the cell wall as part of the APAP1 complex that may be changed as a result of disturbed P4H activity.

## ACKNOWLEDGEMENTS

The authors gratefully acknowledge the financial support by the National Science Center Poland (Sonata16, no 2020/39/D/NZ9/00232). Also, this work has been financed by the European Regional Development Fund of the European Union and Greek national funds through the Operational Competitiveness, Entrepreneurship and Innovation, under the call RESEARCH-CREATE-INNOVATE (project code: T2EDK-01332: n-Tomatomics - Development of new tomato cultivars by using -omics technologies). We would like to thank Mr Emil Zięba for his excellent technical assistance.

## AUTHOR CONTRIBUTION

N.K. performed experiments, interpreted results, prepared data presentation and wrote the first version of the manuscript, A.L. designed research, interpreted results, and corrected the manuscript; project administration, and funding acquisition, A.Z. (Adrian Zając) assisted with the molecular analyses, M.S.C. carried out FTIR analysis, T.S. imaged using SEM and TEM; L.E., D.K., A.K., K.M., EL, P.K. created and validated the cGFP, nGFP transgenic lines and PK revised the manuscript, A.Z. (Artur Zdunek) revised the manuscript.

## DATA AVAILABILITY STATEMENT

The data underlying this article will be shared on reasonable request to the corresponding author.

